# Extracellular matrix scaffold-assisted tumor vaccines induce tumor regression and long-term immune memory

**DOI:** 10.1101/2023.09.12.557449

**Authors:** Sanjay Pal, Rohan Chaudhari, Iris Baurceanu, Brenna J. Hill, Bethany A. Nagy, Matthew T. Wolf

## Abstract

Injectable scaffold delivery is an immune engineering strategy to enhance the efficacy and reliability of cancer vaccine immunotherapy. The composition and structure of the biomaterial scaffold determines both vaccine release kinetics and inherent immune stimulation via the scaffold host response. Extracellular matrix (ECM) scaffolds prepared from decellularized tissues initiate an acute alternative inflammatory response following implantation, which facilitates wound healing following tumor resection and promotes local cancer immune surveillance. However, it remains unknown whether this environment is compatible with generating protective anti-tumor cytotoxic immunity with local immunotherapy delivery. Here, we engineered an ECM scaffold-assisted therapeutic cancer vaccine that maintained an immune microenvironment consistent with tissue reconstruction. Immune adjuvants MPLA, GM-CSF, and CDA were screened in a cancer vaccine formulated for decellularized small intestinal submucosa (SIS) ECM scaffold co-delivery. Though MPLA and GM-CSF showed the greatest increase in local myeloid cell infiltration, we found that the STING pathway adjuvant CDA was the most potent inducer of cytotoxic immunity with SIS-ECM scaffold delivery. Further, CDA did not diminish hallmark ECM immune responses needed in wound healing such as high *Il4* cytokine expression. SIS scaffold delivery enhanced therapeutic vaccine efficacy using CDA and the antigen ovalbumin, curing greater than 50% of established EG.7 tumors in young mice and 75% in 24-week-old mature mice, compared to soluble components alone (0% cured). SIS-ECM scaffold assisted vaccination extended antigen exposure, was dependent on CD8^+^ cytotoxic T cells, and generated long term anti-tumor memory at least 7 months post-vaccination in both young and mature-aged mice. This study shows that an ECM scaffold is a promising delivery vehicle to enhance cancer vaccine efficacy while being orthogonal to characteristics of pro-healing immune hallmarks.

## Introduction

Cancer immunotherapy has been a revolutionary step in cancer treatment. Several classes of immunotherapy have generated curative responses when treating solid tumors including immune checkpoint inhibitors, adoptive T cell therapy, and therapeutic cancer vaccines [1]. However, these successes are not uniform, and tumor regression occurs in only a minority of patients necessitating immune engineering strategies to enhance efficacy and reliability [2]. Biomaterial scaffold-assisted cancer vaccines are one such strategy. Injectable scaffolds from diverse compositions have been engineered to enhance anti-tumor immune responses via material properties that prolong vaccine component delivery and by synergizing with biomaterial induced leukocyte recruitment and activation that occurs during the host immune response [3]. The goal is that the scaffold immune response acts synergistically with an appropriate immune adjuvant to accentuate the density and activity of antigen presenting cells (such as dendritic cells and macrophages) to prime adaptive immune cells. The net result is a coordinated cytotoxic T cell response that is tumor antigen specific and can establish protective immunological memory. This is in contrasts to systemic drug delivery modalities such as nanoparticles where biomaterials are often formulated as a passive vehicle with minimal immune activation.

Extracellular matrix (ECM) scaffolds are medical devices derived from decellularized mammalian tissues that are increasingly used biomaterials in cancer care, where they are implanted in patients during reconstructive surgeries immediately following tumor resection [4–8]. ECM scaffolds are immunomodulatory but generate an immune environment that is entirely distinct from the foreign body reaction generated by many synthetic, polymeric scaffold materials. ECM scaffolds initiate a complex local immune profile that includes a mixed M1/M2 milieu of macrophage phenotypes [9, 10], a Th2 biased CD4+ T helper cell response [11, 12], and Type 2 immune signatures such as eosinophils and elevated IL-4 cytokine [10–12]. ECM scaffold immune modulation has been shown to be indispensable to tissue repair and constructive scaffold remodeling [11, 13–15], and recently, this local ECM scaffold immune environment had been shown to delay tumor formation when seeded with aggressive melanoma cells synergistically with systemic administration of immune checkpoint blockade immunotherapy [10]. Though the local immune environment is inhospitable to tumor progression, cytotoxic effector cells that are needed for treating distant tumors and metastases, such as CD8+ cytotoxic T cells and Natural Killer (NK) cells, were only nominally enriched in this ECM immune environment. While several studies have investigated biomaterial assisted cancer vaccines composed of polymer scaffolds[16–21], few have examined the potential of ECM scaffolds in cancer vaccine delivery [22–24].

Therefore, a critical question is whether the ECM scaffold immune environment can synergize with local cancer immunotherapy strategies to engage tumor specific cytotoxic adaptive immune cells (e.g., cytotoxic T cells) while also preserving the beneficial pro-regenerative hallmarks of the ECM scaffold response. The objectives of the present study were to leverage the material and immune modulating properties of ECM scaffolds to develop an effective therapeutic cancer vaccine, and for this formulation be orthogonal to ECM immune biomarkers that had been previously associated with successful tissue reconstruction. To achieve this goal, we first characterized the local, regional, and systemic immunological changes when ECM scaffolds are used to deliver immune stimulating adjuvants. We then utilized an *in vivo* cytotoxic lymphocyte assay to elucidate an adjuvant formulation that induced antigen-specific cellular immunity when delivered with an ECM scaffold and characterized antigen release properties. Finally, we implemented this formulation to determine feasibility of a biologic scaffold vaccine to therapeutically treat established tumors and to decipher the immune mechanisms of this response.

## Results

### The ECM scaffold immune microenvironment is temporally modified by immune adjuvant co-delivery and depends on adjuvant type

Decellularized porcine small intestinal submucosa (SIS) ECM was prepared for use as an archetypical ECM scaffold biomaterial and cryogenically milled into an injectable particulate (**Fig 1A**). SIS was selected because it has been extensively studied in soft tissue repair in both pre-clinical and clinical models, including in cancer treatment [7, 8, 25, 26]. An injectable formulation permits minimally invasive localized delivery and avoids the trauma of surgical implantation thus isolating the host response to the ECM scaffold itself (**Fig 1B**). Decellularization was confirmed by lack of visible nuclei with H&E and DAPI staining of histologic sections (**Fig 1C, D**) and a 95% reduction in double stranded DNA content (360.7 ng/mg in SIS vs 6,861.1 ng/mg in native intestine, dry weight, **Fig 1E**). The resulting SIS particles ranged between 41-70 µm in size as quantified from scanning electron microscopy (SEM) images (95% confidence interval), while still preserving characteristic ECM structures such as the banding pattern found in intact triple helical collagen fibrils (**Fig 1F, G**). SIS particles were sterilized via ionizing irradiation and pathogen screened prior to *in vivo* studies.

**Figure 1:**
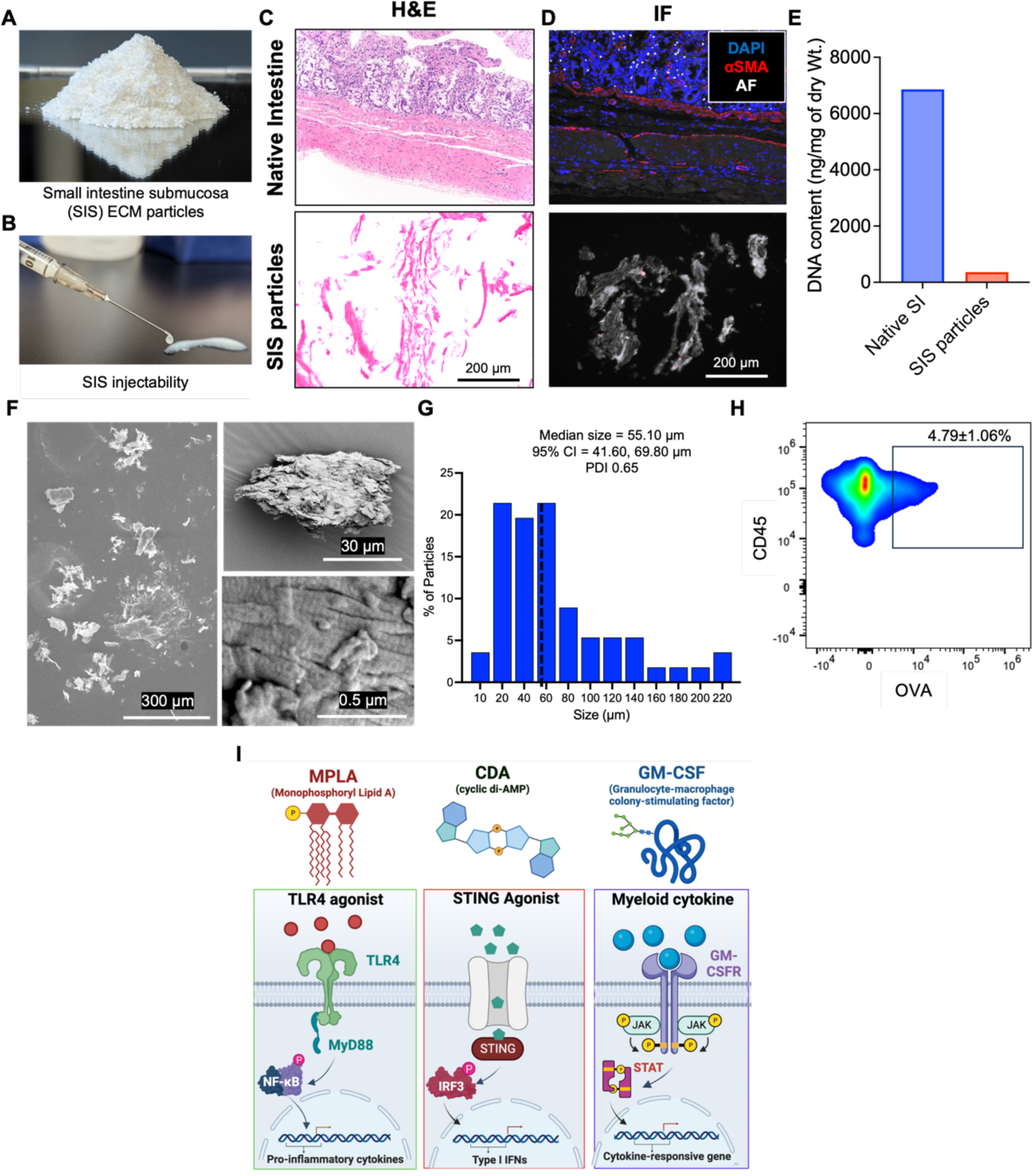
Injectable SIS ECM particle characterization and immune adjuvant selection. (A) SIS particles prepared from cryogenically milled decellularized porcine submucosa, which (B) was hydrated for injectable delivery. SIS decellularization compared to native intestine was confirmed via (C) H&E histology of SIS particles, (D) removal of nuclei (DAPI) and smooth muscle actin (aSMA) staining, and (E) reduction of total double stranded DNA content via the PicoGreen assay. (F) SEM imaging of SIS particle topography and (G) particle size distribution. (H) Quantification of ovalbumin antigen (OVA) uptake in immune cells infiltrating SIS scaffolds 3 days after subcutaneous implantation in mice. (I) Mechanism of action of SIS particles combined with one of three immune adjuvants for subsequent studies: MPLA, CDA, and GM-CSF (Created with BioRender.com). AF, autofluorescence; SI, small intestine; CI, confidence interval; PDI, polydispersity index.

Tumor antigen uptake is the first step of generating cytotoxic cellular immunity during cancer vaccination, and since previous studies showed that leukocytes readily infiltrated ECM scaffold biomaterials [10, 11, 13, 27], we sought to determine if these cells were capable of internalizing exogenous vaccine antigen. We adsorbed the model antigen ovalbumin (OVA) to SIS ECM for subcutaneous injection in mice and found uptake in nearly 5% of infiltrating CD45^+^ immune cells within 3 days (**Fig 1H**). We then investigated whether immune adjuvant co-delivery could activate antigen specific cytotoxic immunity against this delivered antigen within the SIS ECM immune microenvironment without diminishing other immune hallmarks associated with healing. We selected adjuvants that stimulate minimally overlapping immune activation pathways and have shown promise in clinical vaccine trials. MPLA (Monophosphoryl Lipid A) is a TLR-4 agonist transduced through the MyD88 and TRIF signaling pathways [28, 29]; CDA (the cyclic di-AMP analog 2’3’-c-di-AM(PS)2 (Rp,Rp)) which activates the STING pathway to produce type I interferons [30–32]; GM-CSF (Granulocyte-macrophage colony-stimulating factor) which mobilizes myeloid cells and promotes differentiation to antigen presenting cells (**Fig 1I**) [33, 34]. To preserve native ECM scaffold properties, we did not modify ECM composition and used passive adsorption and absorption of each adjuvant with lyophilized SIS ECM particles prior to subcutaneous injection in C57Bl/6 mouse flanks.

We first characterized how each adjuvant modulated the local SIS ECM scaffold immune microenvironment and found profound changes to both leukocyte recruitment and spatial distribution. During the acute phase of the immune response, SIS ECM particles aggregated at the injection site with acute host cell infiltration concentrated around the implant border with sporadic clusters of cell accumulation within inter particle spaces (**Fig 2A arrows**). In the absence of adjuvant, total cellularity modestly peaked at 7 days (3,221 cell/mm^2^), but remained relatively constant over the 1-14 day time course (**Fig 2A,B**). The TLR4 agonist MPLA caused the most striking increase on SIS cellularity of all adjuvants tested, recruiting nearly 4,896 cells/mm^2^ after 1 day that remained chronically elevated at this level after 14 days creating an infection-mimicking appearance with abundant polymorphonuclear cells (**Fig 2A, B**). The cytokine GM-CSF also rapidly induced high acute inflammatory response as MPLA after 1 day similar to MPLA but had returned to SIS control levels by day 14. Unexpectedly, the STING agonist CDA decreased initial cell recruitment by approximately 20% after 1 day before returning to control SIS levels. Though fewer cells were observed, there was no evidence of local tissue necrosis in adjacent muscle, adipose, or hypodermis with CDA delivery, nor with other adjuvants. STING activation-induced apoptosis has been observed in specific cell populations such as B cells after chronic exposure, but not in fibroblasts [35], however the decreased cellularity in this study appears to be short lived. Spatially, cellularity was greatest at the SIS interface (within 200 µm of the implant border) with relatively few cells infiltrating the core. Evidence of ECM degradation via fragmentation was most prominent with MPLA co-delivery where it increased the aggressiveness of this remodeling response (**Fig 2A**).

**Figure 2.**
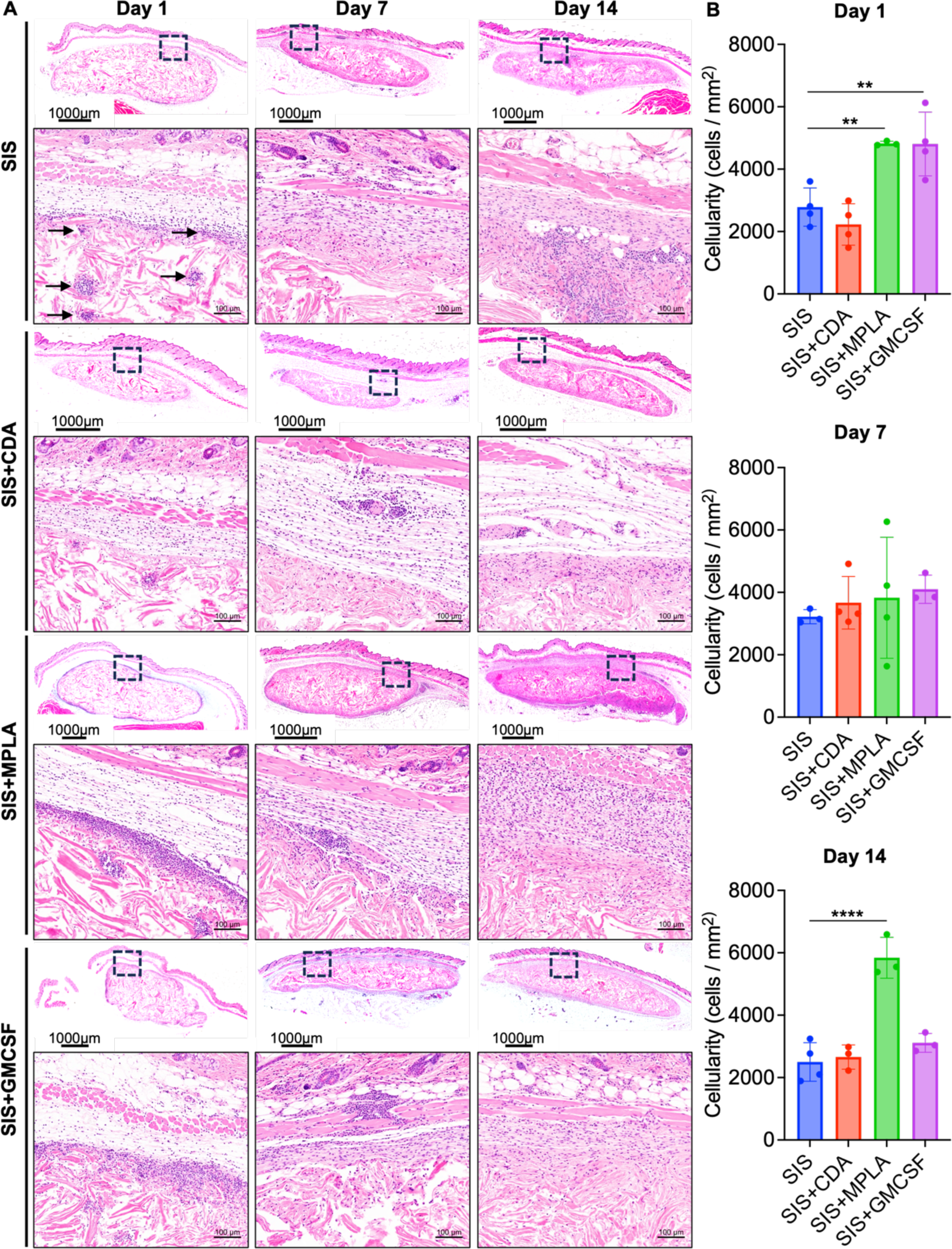
Histologic host response to SIS ECM implantation with immune adjuvant co-delivery. (A) H&E stained images showing the morphology of subcutaneously injected SIS ECM particles in C57Bl/6 mice alone or with the immune adjuvants CDA, MPLA or GM-CSF at 1, 7 and 14 days post implantation. Boxes highlight immune infiltrates at the scaffold interface (20X). (B) Total cell density was quantified at the SIS ECM interface across the entire section (N=3-4, mean ± SD). \*\**p* < 0.01, \*\*\*\**p* < 0.0001, one-way ANOVA with Dunnett’s multiple comparisons test.

Myeloid lineage cells of the innate immune system are the first responders to implanted biomaterials, play a deterministic role in downstream scaffold remodeling [13], and are crucial to the vaccine response [16, 18]. We therefore sought to determine the immune phenotype and distribution of SIS ECM infiltrating cells by multiplex immunofluorescence staining. Neutrophils (Ly6G^+^), macrophages (F4/80^+^), and antigen presenting cells (APCs, CD86^+^) were selected to characterize the myeloid cell types that define the host immune response to biomaterials and antigen presentation for vaccines (**SFig 1B**). Neutrophils and APCs accounted for the majority of myeloid cells in the acute SIS ECM control response 1 day post-implantation and then completely subsided, giving way to macrophages by day 7 at the SIS interphase (**Fig 3A,B and SFig 1A**). Adjuvant co-delivery increased neutrophil infiltration 2-fold in MPLA and GM-CSF groups, compared to SIS controls at the interphase (**Fig 3A,B**). CDA delivery did not alter neutrophil density, though slight decreases in other populations increased overall proportion (**Fig 3C**). Myeloid infiltration was substantially lower in the implant core for each group and consisted primarily of neutrophils after 1 day that was significantly increased by GM-CSF delivery (**Fig 3D**). For each group, macrophage infiltration within the core gradually increased to 14 days, and APCs remained rare (**Fig 3D and SFig 1C**).

**Figure 3.**
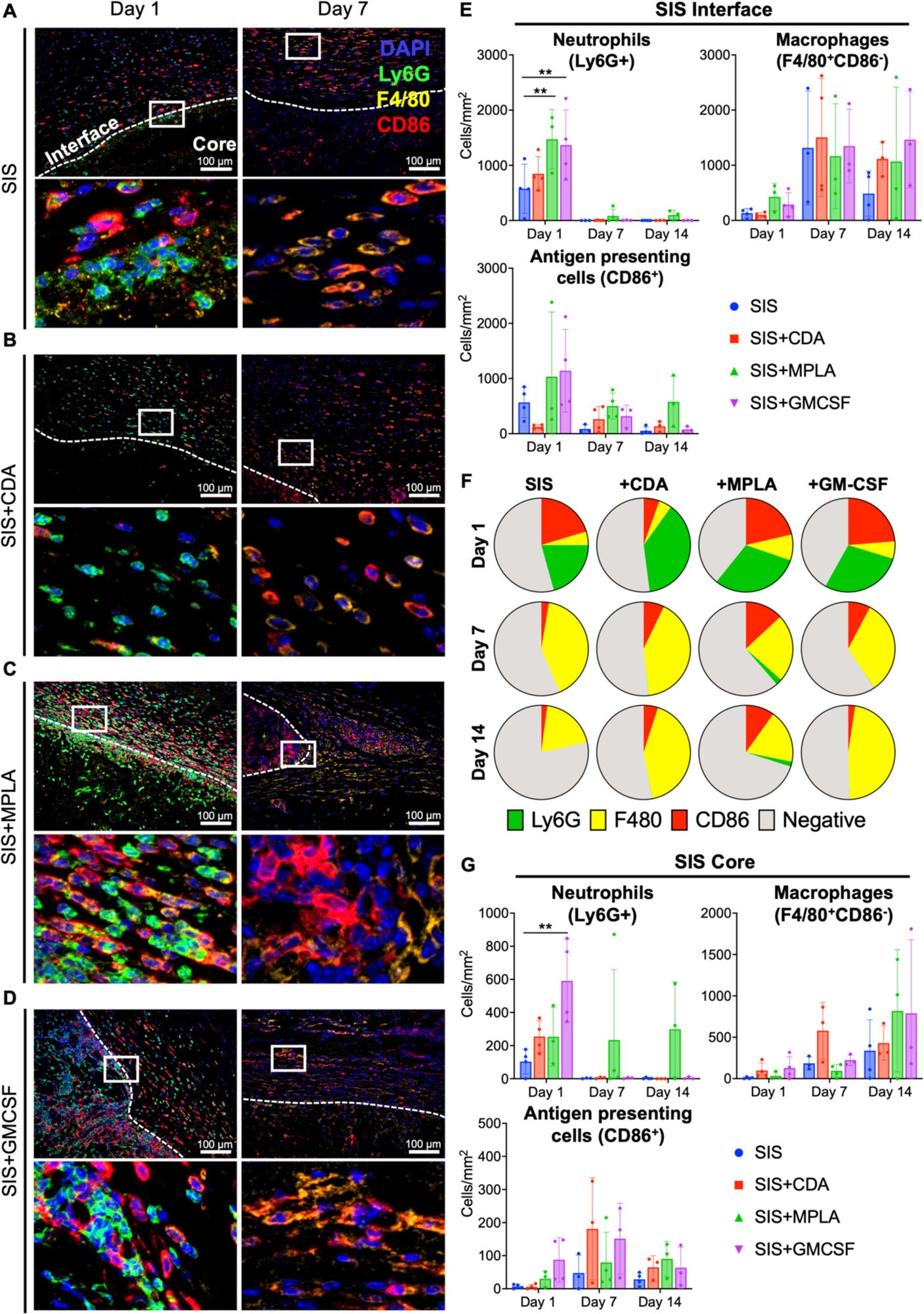
Spatiotemporal immune profiling SIS ECM scaffolds co-delivered with immune adjuvant. (A-D) Multiplex immunofluorescent images of subcutaneously injected SIS ECM particles in C57Bl/6 mice alone or with the immune adjuvants CDA, MPLA or GM-CSF at 1 and 7 days post implantation (20X objective). Day 14 images are available in SFig 1C. Dashed lines delineate the SIS ECM implant border and boxes detail immune phenotype at this interface. The interface was defined as 200 um concentrically from the border, and the core as greater than 200 um towards the center SFig A. (E) Myeloid cell density (N=3-4, mean ± SD) and (F) proportions of each cell type quantified across the entire SIS interface. (G) Myeloid cell density quantification within the SIS ECM implant core with adjuvant (N=3-4, mean ± SD). \*\**p* < 0.01, two-way ANOVA with Tukey’s multiple comparisons test.

The local immune response to a biomaterial scaffold is primarily composed of myeloid cell infiltration, however, antigen specific lymphocyte priming most efficiently occurs in lymphoid tissues such as regional draining lymph nodes. Previous studies have shown that IL-4 expression is a hallmark of the pro-regenerative ECM scaffold immune environment [10–12], thus we used *Il4* gene expression in SIS implant draining lymph nodes as an indicator of regional immune modulation by ECM and to determine whether this response is perturbed by adjuvant. Conversely, *Ifny* (encoding cytokine interferon-gamma) expression is a biomarker of Th1 immunity that can be induced by certain cytotoxic inducing adjuvants and during infection. Since adaptive priming takes several days to develop, we evaluated lymph nodes 7- and 14-days post implantation, compared to a naïve control (without ECM or adjuvant injection). SIS ECM alone induced 7.5-fold increased lymph node *Il4* expression compared to naïve lymph nodes, and unexpectedly retained elevated expression in all adjuvant groups, and both MPLA and GM-CSF significantly increased expression relative to SIS only controls at 7 days. Most adjuvant groups returned to SIS control Il4 expression baseline by 14 days except GMCSF which showed a 9-fold decrease (**Fig 4A**). The Th1 associated gene Ifny showed a slight decrease with ECM implantation compared to naïve mice, with no significant modulation with adjuvant (**Fig 4B**). These results show that adjuvant delivery is largely orthogonal to Th2 activation features of the ECM host response.

**Figure 4.**
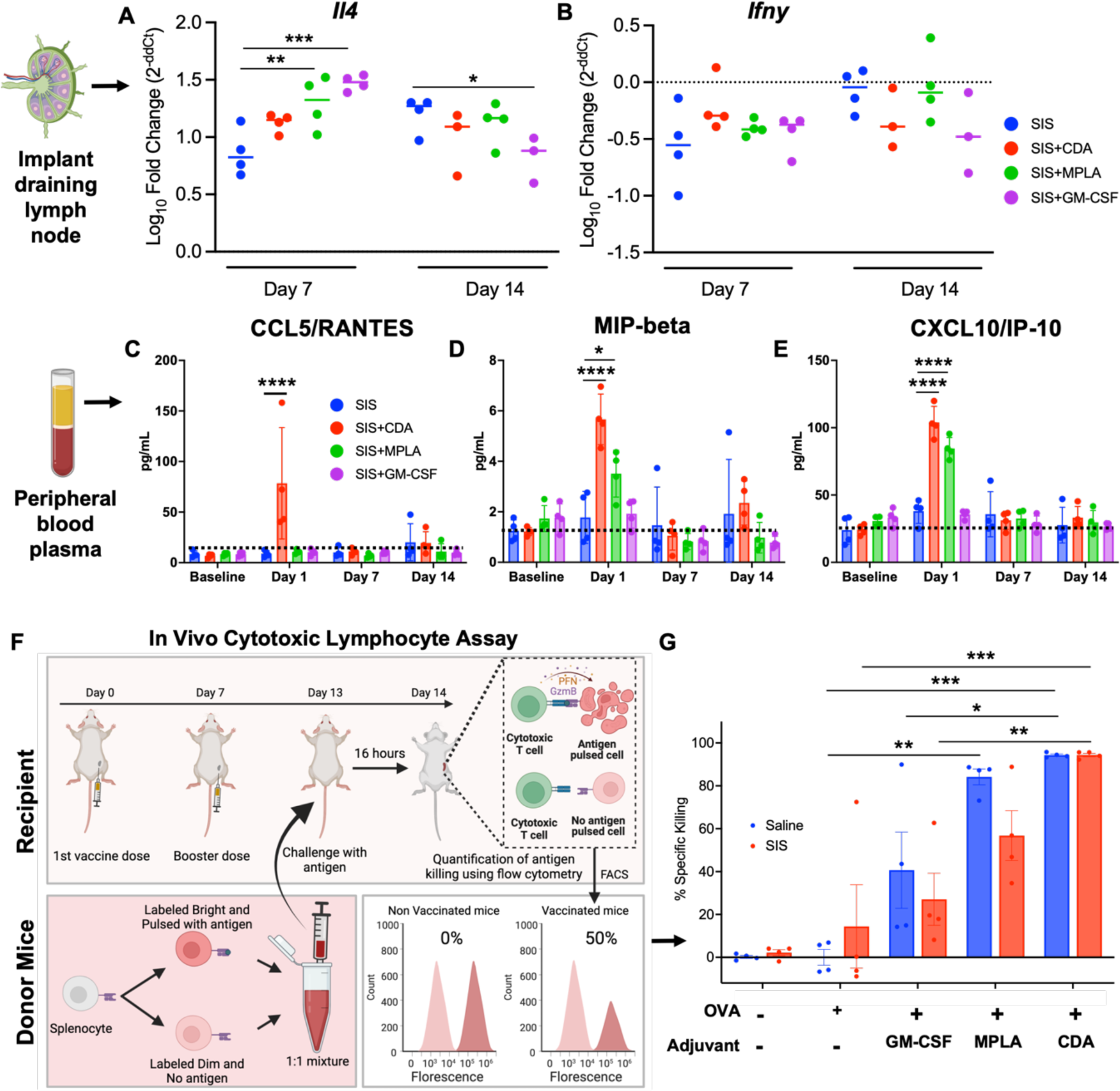
Lymph node and systemic immune modulation with ECM and adjuvant delivery. Quantitative real time PCR of (A) *Il4* and (B) *Ifny* gene expression in scaffold draining lymph nodes 7 and 14 days after SIS ECM co-delivery with adjuvants. (N=3-4, mean ± SD). \*\**p* < 0.01, \*\*\**p* < 0.001, two-way ANOVA with Šídák’s multiple comparisons test (C-E) Peripheral blood was collected 2 days before SIS ECM implantation (Baseline) then 1, 7, and 14 days post implantation for Luminex analysis. (N=3-4, mean ± SD). \**p* < 0.05, \*\*\*\**p* < 0.0001, two-way ANOVA with Tukey’s multiple comparisons test. (F) Schematic of the in vivo cytotoxic lymphocyte assay procedure (Created with BioRender.com). (G) Quantification of OVA antigen-specific cytotoxic T cell killing when SIS ECM was co-delivered with each adjuvant type. (N=3-4, mean ± SD). \**p* < 0.05, \*\**p* < 0.01, \*\*\**p* < 0.001, two-way ANOVA with Tukey’s multiple comparisons test.

An advantage of biomaterial vaccine delivery strategies is the ability to achieve high local concentrations that would otherwise be toxic if given systemically. We analyzed peripheral blood plasma using a Luminex cytokine panel 1, 7, and 14 days after adjuvant co-delivery with SIS-ECM to evaluate risk of deleterious systemic dysregulation, such as cytokine storm. SIS ECM implantation alone did not affect circulating cytokine levels in any of the 21 analytes tested (**Fig 4C**, **Supplementary Tables 2**). CDA and MPLA adjuvant co-delivery induced only transient elevation of specific cytokines and chemokines 1 day after implantation resolving to baseline by 7 days (**Fig 4C-E and SFig 2A**). CCL5 and MIP-1 beta were increased by CDA delivery, and IP-10 increased with either CDA or MPLA adjuvant delivery (**Fig 4C-E**). These results show that adjuvant delivery with ECM is tightly regulated with only acute perturbations to homeostasis that are not long lasting.

### The STING agonist CDA optimally induces antigen-specific cytotoxic T cell activity in the ECM immune environment

The primary objective of a cancer vaccine is to prime and activate antigen specific anti-tumor lymphocytes to systemically target neoplasms. We performed an *in vivo* cytotoxic T lymphocyte (CTL) assay to quantify T cell activity to determine whether the ECM host immune microenvironment was compatible with vaccination using the tested adjuvants using the antigen ovalbumin (OVA). Mice were administered a subcutaneous flank priming and booster vaccination dose consisting of SIS, adjuvant, and OVA at day 0 and day 7, respectively, and challenged with adoptively transferred splenocytes pulsed with either the MHCI (H-2Kb) restricted OVA peptide SIINFEKL or negative control (**Fig 4F**). Vaccine efficacy was calculated by antigen-specific killing efficiency of transferred cells harvested from systemic lymphoid tissue, spleen. As expected, SIS ECM alone did not reliably induce OVA specific immunity and required cytotoxic-inducing adjuvant. GM-CSF was the least effective adjuvant for both ECM and Saline controls. MPLA generated 82% cell killing when delivered as a soluble vaccine, but was less effective and consistent with SIS delivery at 50% killing. In contrast, CDA was the most effective cytotoxic adjuvant with ECM delivery, generating greater than 95% specific killing with both Saline and ECM (**Fig 4G**). We then performed CDA titrations between 0.2-20 µg using a simplified version of the CTL assay and found similar cytotoxic activity when delivered with SIS ECM or Saline at all concentrations, including a small decrease in potency at the lowest dose of 0.2 µg (**SFig 2B**). This suggests the mechanism of action for the STING agonist CDA is compatible with vaccination when delivered with an ECM scaffold biomaterial.

### ECM scaffolds prolong antigen retention and release kinetics

A key feature of biomaterial scaffold vaccine delivery is the ability to modify retention and exposure of vaccine components, and the optimal delivery kinetics likely varies empirically by specific formulation and mechanism. We characterized retention and release kinetics of vaccine components from SIS ECM using fluorescent live animal imaging following a single subcutaneous vaccine dose including either fluorescently labeled OVA protein or a CDA analogue (cGAMP, cyclic guanosine–adenosine monophosphate) [36]. In addition to soluble delivery with Saline, the inorganic vaccine adjuvant aluminum hydroxide (Alum) was used as a positive control due to its established effects as a depot for protein adsorption and widespread clinical usage [37]. SIS ECM extended the retention and release of the whole-protein antigen OVA. After a 2-3 day burst release phase, OVA signal slowly decreased to background over 30 days with only 50% release after 4 days (**Fig 5A,B and SFig 3A, B**). In comparison, soluble OVA protein was rapidly cleared from the injection site with almost 80% release after 4 days (**Fig 5A,B and SFig 3A, B**). The positive control Alum showed the slowest release with only 20% release after 4 days (**Fig 5A,B and SFig 3A, B**). These same trends were maintained with or without CDA delivery with OVA (**SFig 3A, B**). Conversely, release kinetics of the CDA analogue cGAMP was substantially more rapid than OVA in all groups, with total clearance (greater than 85%) within 4 days of soluble delivery with Saline (**Fig 5C,D and SFig 3C**). ECM did not significantly alter cGAMP release kinetics, though Alum did prolong release with only ∼50% release after 4 days (**Fig 5C, D and SFig 3C**).

**Figure 5.**
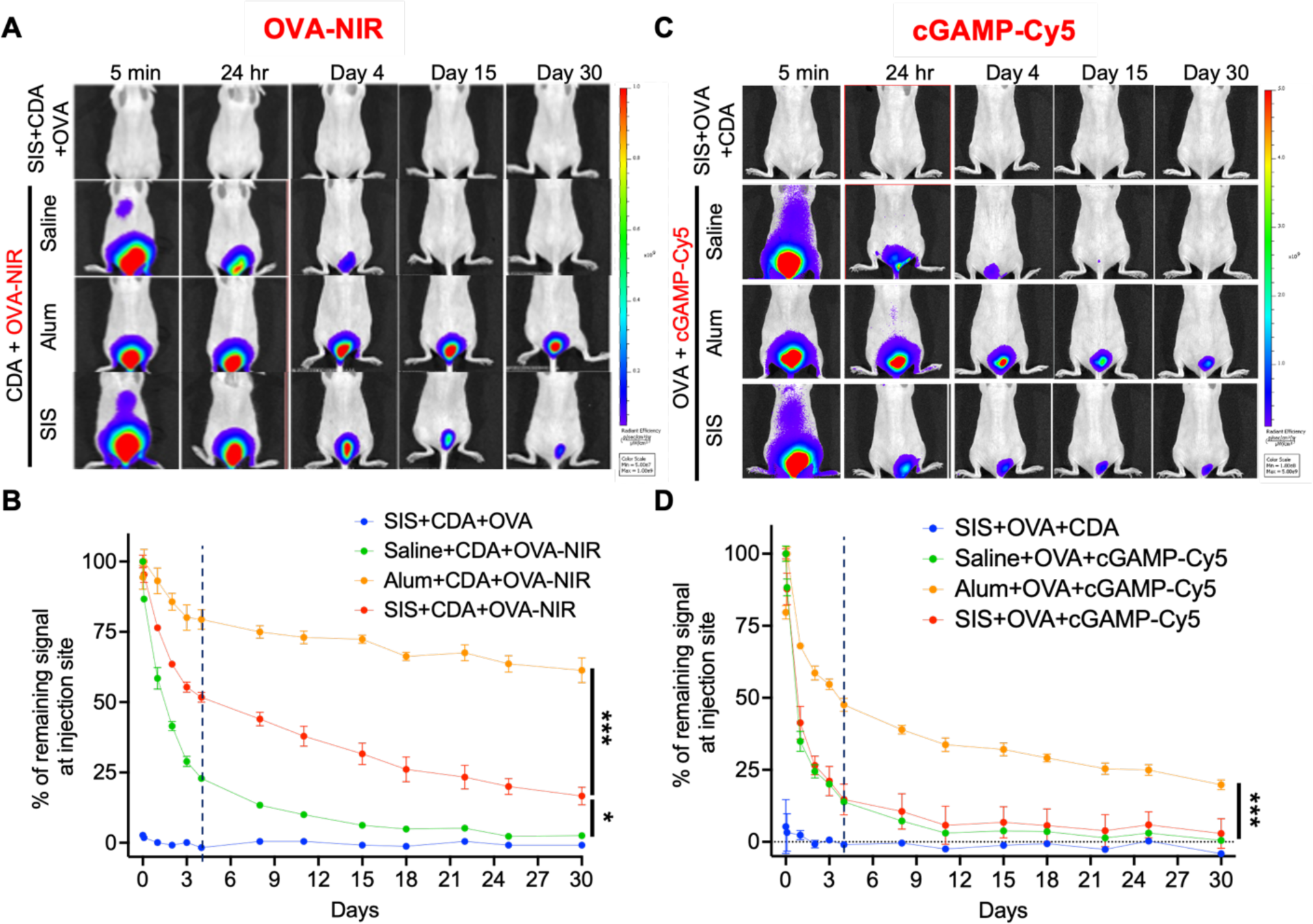
In vivo vaccine release kinetics from SIS scaffolds via live animal fluorescence imaging. (A) Antigen retention was tracked using Licor800 NIR dye conjugated OVA protein when co-delivered subcutaneously with CDA and either SIS ECM, Alum, or Saline control at the tail base of hairless immunocompetent SKH1 mice. (B) Fluorescence flux from labeled OVA was quantified at the tail base and normalized to initial signal (5 min post injection) over 30 days. (N=3, mean ± SD) (C) Cyclic dinucleotide retention was tracked using a Cy5 conjugate of the CDA analogue cGAMP co-delivered with unlabeled OVA and each biomaterial condition. (D) Fluorescence flux from labeled cGAMP was quantified at the tail base and normalized to initial signal (5 min post injection) over 30 days. (N=3, mean ± SD). \**p* < 0.05, \*\*\**p* < 0.001, two-way ANOVA with Tukey’s multiple comparisons test.

### An ECM scaffold assisted therapeutic cancer vaccine induces CD8+ T cell dependent tumor regression and protective anti-tumor immune memory

We tested the efficacy of an ECM scaffold assisted vaccine to treat established tumors. Ovalbumin expressing EG.7-OVA mouse lymphoma cells were injected subcutaneously in the flanks of both young 8-week-old and mature 24-week-old C57Bl/6 mice and treated once the tumors grew to 75-100 mm^3^ and again 7 days later (**Fig 6A**). An ECM assisted cancer vaccine was subcutaneously injected caudally to the tumor site using the same formulation used in the CTL assay (5 mg SIS, 20 µg CDA, 100 µg OVA), using Alum as a reference particulate material. A therapeutic SIS ECM scaffold assisted cancer vaccine induced curative lymphoma tumor regression in 57% of young mice (4/7), whereas soluble OVA and CDA delivery in Saline controls did not produce complete responses in any treated animals (0/6) (**Fig 6B, C, E**). Vaccination with Alum was similar to ECM with 50% regression (3/6) confirming the importance of material delivery to enhance therapeutic efficacy (**Fig 6B,C,E**). Tumor growth kinetics showed that SIS ECM delivery quickly induced tumor regression to undetectable sizes in all animals, though nearly half underwent local recurrence within 1 week of the second dose (**Fig 6B,D**). Saline delivery induced some tumor regression or growth stasis in most animals, and all experienced recurrence (**Fig 6B, D**). We also confirmed that CDA was necessary as an immune adjuvant as OVA delivery alone with ECM or Alum did not show any regression in the tumor growth kinetics (**SFig 4A,B**). Young immunologically naïve mice are useful models for immunology research, however, are still rapidly developing and do not reflect the macroenvironment of adults in which most cancers arise. Since age is a crucial variable in immunotherapy responsiveness [38], we tested SIS ECM scaffold vaccine efficacy in mature adult mice (24 weeks old) (**Fig 6F**). We found that SIS ECM scaffold vaccines were highly effective in mature mice, with 77% durable tumor regression (7/9 mice) (**Fig 6G,H**). Recurrence occurred approximately 3 weeks after the second treatment dose. These results show that ECM can be used to enhance efficacy of a therapeutic vaccine efficacy in both young and mature adult mice.

**Figure 6.**
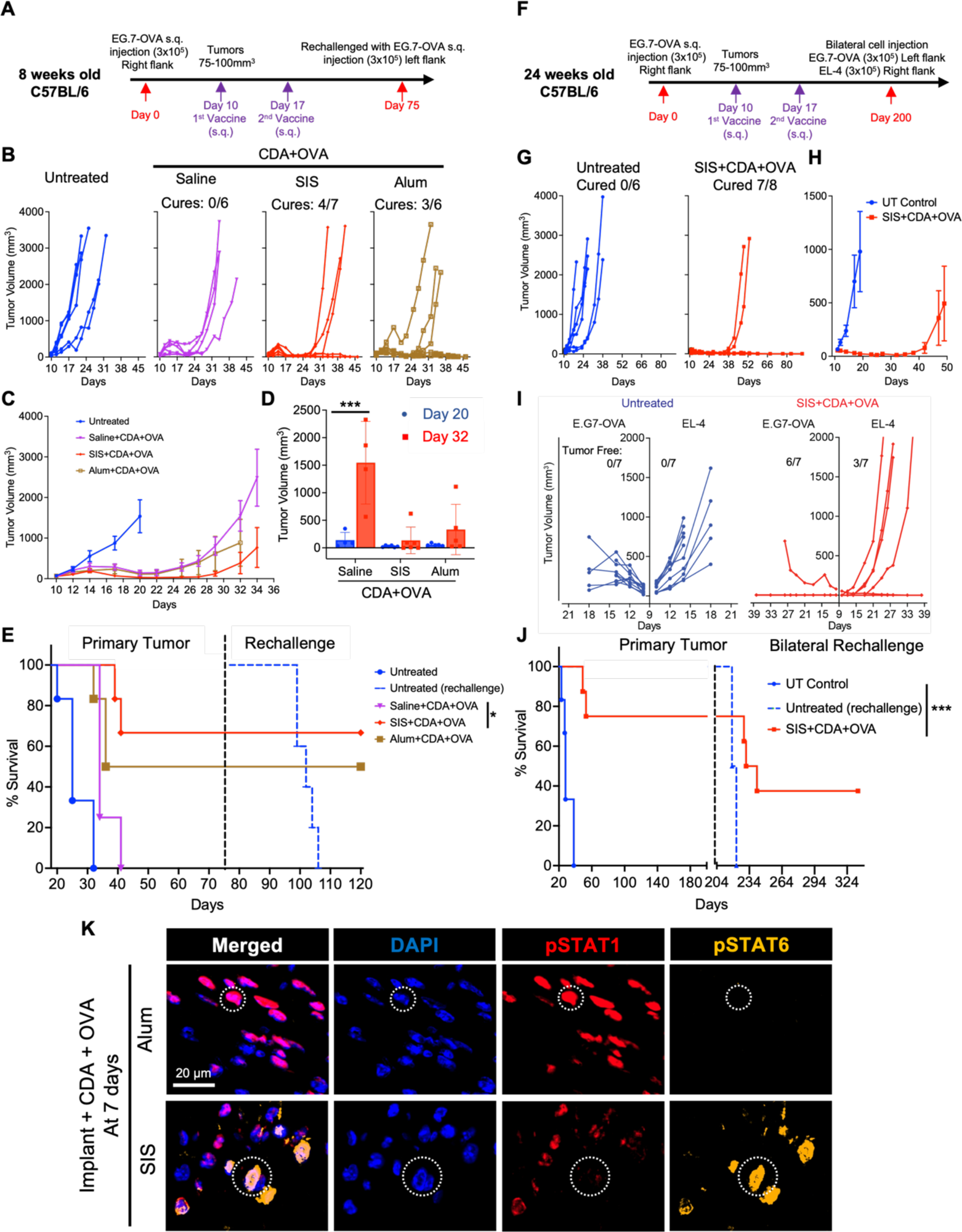
Tumor regression and long-term protection following therapeutic SIS ECM scaffold assisted vaccination of established tumors in young and adult-mature mice. (A, F) Schematic timeline of E.G7-OVA tumor induction, subcutaneous vaccination schedule using CDA and OVA antigen, and tumor rechallenge in 8-week-old young and 24-week old adult mature C57Bl/6 mice. (B) Individual tumor growth curves for scaffold assisted therapeutic vaccination, with cures defined as complete and durable tumor regression to a minimum of 75 days. (C) Average tumor growth kinetics and (D) quantification 20 and 32 days after tumor induction as early and late responses. (N=6-7, mean ± SEM). \*\*\**p* < 0.001, two-way ANOVA with Šídák’s multiple comparisons test. (E) Impact of vaccination on overall survival and on rechallenge with EG.7-OVA cells on the contralateral flank. Log-rank (Mantel-Cox) test. (G) Individual tumor growth curves and (H) average tumor growth kinetics for scaffold assisted therapeutic vaccination, with cures defined as complete and durable tumor regression to a minimum of 200 days. (I) Individual tumor growth curves for scaffold assisted therapeutic vaccination upon bilateral rechallenge of E.G-OVA (Left flank) and EL-4 (Right flank) in primary tumor surviving mice. (J) Impact of vaccination on overall survival and on Bilateral rechallenge with EG.7-OVA and EL-4 cells. (N=6-8, mean ± SEM), Log-rank (Mantel-Cox) test. (K) Multiplex immunofluorescent images showing phospho-STAT1 and phospho-STAT6 staining of subcutaneously injected SIS-Vax and Alum-Vax in C57Bl/6 mice 7 days post implantation (20X objective).

We performed a series of tumor rechallenge experiments to determine anti-tumor immunological memory generation in surviving mice exhibiting durable tumor regression. Young-vaccinated mice were rechallenged 75 days later with EG.7-OVA cells on the contralateral flank relative to the original tumor. All surviving mice from the SIS ECM and Alum vaccine treated groups were protected on rechallenge suggesting systemic immunological memory (**Fig 6E**). We performed a similar experiment in surviving mature-vaccinated mice but used a more aggressive bilateral rechallenge over 200 days post-implantation: EG.7-OVA tumor cells on the right contralateral flank and its parental lymphoma line EL-4 on the ipsilateral flank (**Fig 6F**). The goal of this rechallenge was to determine whether immunological memory persists long term (6 months) and to evaluate potential “epitope spreading” against non-OVA tumor antigens. Since EL-4 is the parental strain from which EG.7-OVA was derived, it shares a similar antigen landscape but without OVA protein. Both tumor lines rapidly grew in age-matched untreated control mice (**Fig 6I**). SIS ECM vaccinated mature mice strongly rejected EG.7-OVA lymphoma cells in 85% of mice (6/7), demonstrating immune protection, and surprisingly, moderate protection against the non-OVA expressing EL-4 line in 42% of mice (3/7) (**Fig 6I**). This data suggests epitope spreading may be a compatible mechanism when vaccines are co-delivered with an ECM scaffold biomaterial. Young-vaccinated mice underwent a similar bilateral rechallenge (**SFig 4C, D**) over 150 days after the first rechallenge. All mice in each of the SIS ECM (N=4) and Alum (N=3) vaccine groups were protected from EG.7-OVA cells, however, only 1 Alum vaccine mouse demonstrated protection against EL-4 (**SFig 4E**). Similar tumor cure rates and long-term memory was observed for both SIS ECM and Alum despite their disparate compositions. We compared the local cytokine signaling milieu between materials to characterize differences in their immune environments and to determine if these were preserved in the complete vaccine composition (both CDA and OVA). We found that vaccine co-delivery triggered STAT1 phosphorylation in the SIS ECM microenvironment (induced by interferon signaling) after 7 days (**Fig 6K**), comingled with cells exhibiting STAT6 phosphorylation (induced by IL-4 signaling) induced by SIS ECM alone (**SFig 4F**). In contrast, cells responding to Alum showed STAT1 activation with rare STAT6 phosphorylated cells. These show that local IL-4 signaling is compatible with cytotoxic immune generation and that SIS ECM and Alum generate different immune environments to produce a similar functional outcome.

After demonstrating efficacy of an ECM therapeutic cancer vaccine, we then sought insights into the mechanism of tumor regression and immune protection, and to determine whether functional cytotoxic anti-tumor immunity was indeed being generated by ECM scaffold vaccine delivery. Cytotoxic effector cells were depleted via intermittent systemic administration of monoclonal antibodies beginning prior to therapeutic vaccination in EG.7-OVA bearing mice and maintained throughout the experiment: anti-CD8b (cytotoxic T cells), anti-NK1.1 (cytotoxic NK cells), isotype negative controls (IgG1), or untreated (**Fig 7A**). Depletion efficiency was greater than 95% for cytotoxic T cells and 81% for NK cells in peripheral blood and the spleen, and depletion alone did not affect primary tumor growth (**Fig 7B, SFig 5A-D**). Tumor regression by SIS ECM vaccine delivery was completely abrogated with CD8b targeted depletion (0/8 cured) showing that cytotoxic T cells are essential effectors (**Fig 7C-F**). NK cell depletion variably influenced vaccine with loss of potency in 25% of mice (3/8 cured) in comparison to isotype control (5/8 cured) (**Fig 7C-F**). This suggests that both CD8 and NK cells contribute to anti-tumor immunity, and that CD8+ cytotoxic T cells are the more essential and potent effectors for tumor control with ECM vaccine delivery.

**Figure 7.**
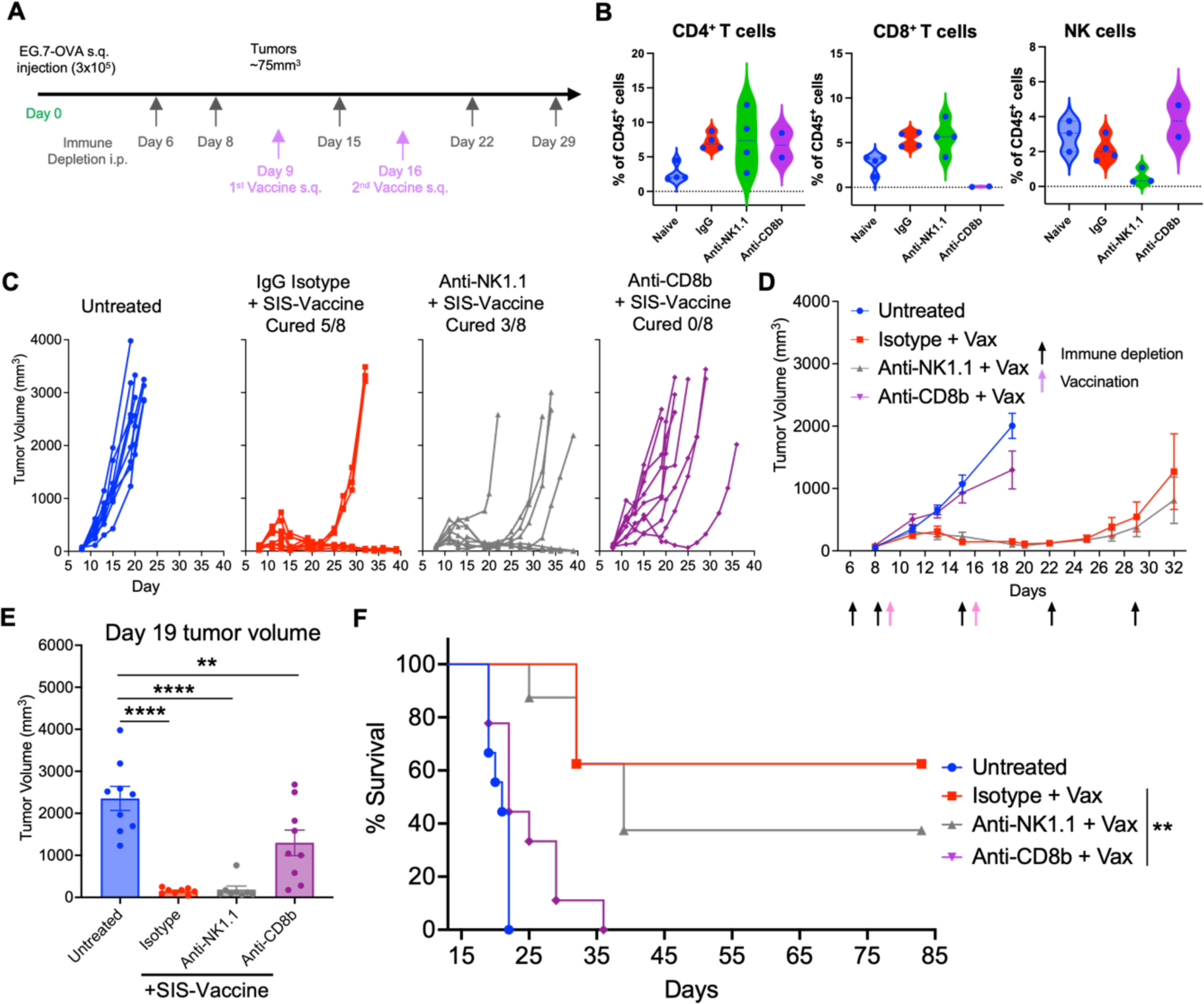
ECM vaccine efficacy following antibody mediated cytotoxic effector cell depletion. (A) Schematic timeline of EG.7-OVA tumor induction, antibody mediated cell depletion schedule, and vaccination schedule using CDA and OVA with SIS ECM. (B) CD8 cytotoxic T cell and NK cell depletion was verified with flow cytometry analysis of peripheral blood. (N=2-4) (C) Individual tumor growth curves with SIS ECM vaccination following each depletion condition, (D) average tumor growth, and (E) tumor volume at 19 days and (F) survival analysis. (N=8, mean ± SEM). \**p* < 0.05, \*\**p* < 0.01, \*\*\*\**p* < 0.0001, one-way ANOVA with Tukey’s multiple comparisons test. Survival comparison Log-rank (Mantel-Cox) test.

## Discussion

We found that injectable decellularized ECM scaffolds enhance therapeutic cancer vaccine efficacy when combined with the appropriate immune adjuvant. ECM scaffolds infused with tumor protein antigen and the STING agonist CDA enhanced antigen-specific cytotoxic T cell immunity, induced curative regression of established tumors, and generated protective anti-tumor memory. ECM scaffolds were inherently immune modulatory, locally recruiting macrophages and antigen presenting cells to the local vaccination site, and prolonged antigen retention. With the addition of immune adjuvant this inflammation remained localized and resolved without adverse events such as systemic toxicity, autoimmunity, or local tissue damage demonstrating safety. Further, cytotoxic immune activation against tumor antigen were orthogonal to IL-4 cytokine signaling elicited by ECM scaffold materials as hallmarks of the pro-regenerative immune response. These results show that alternative forms of biomaterial inflammation are conducive to cytotoxic targeting immunotherapy and is not limited to synthetic scaffolds that stimulate inflammation characteristic of the foreign body reaction.

Local leukocyte recruitment to injectable scaffolds carrying cancer vaccine is an important mechanism for augmenting immune recognition. We confirmed that SIS ECM particles triggered a robust immune infiltrate at the vaccination site, and that these cells were capable of internalizing exogenous antigen (**Fig 1H**). Total cell densities in response to SIS ECM were qualitatively similar to previous reports of subcutaneously implanted ECM [10, 13, 39], with a peak in macrophage recruitment within 1 week of implantation. Macrophages are critical mediators of ECM scaffold remodeling during tissue repair and are among the most well-studied cell type in the ECM host response, though we identified additional vaccine-relevant dynamics that had not been previously described. Of interest were APCs, which are required for vaccine antigen uptake and subsequent T cell priming via co-stimulatory ligands such as CD86. Surprisingly, APC recruitment to SIS ECM scaffolds peaked early (1 day post implantation) and had decreased substantially by 7 and 14 days, which closely followed Ly6G^+^ neutrophil dynamics. Additionally, we found instances of intracellular Ly6G within APCs suggesting neutrophil efferocytosis (clearance of apoptotic cells). Previous studies showed that neutrophils were not essential to ECM scaffold mediated muscle repair [27], though efferocytosis can contribute to downstream T cell function during infection [40]. SIS ECM displayed an adjuvant-like effect by recruiting numerous immune cells however, this immune response alone did not consistently generate functional cytotoxic T cell responses in a CTL assay (**Fig 4G**), which motivated co-delivery with exogenous adjuvants to stimulate this branch of adaptive immunity.

We subsequently identified adjuvant interactions with the SIS ECM immune response that informed optimal vaccine design. Combining scaffold with adjuvant is ideally synergistic wherein the scaffold immune response attracts leukocytes that are then stimulated by locally high concentrations of immune adjuvant. Since the duration, intensity, and phenotype of immune response varies with scaffold composition, we hypothesized that the optimal adjuvant varies with biomaterial type. This approach has identified promising candidates for alginate cyrogels, inorganic silica rods, porous polyesters, and hydrogels [17, 18, 41–43], though there has been limited investigation in ECM biomaterials that create a unique and disparate immune Type 2-biased environment. We found that the TLR4 agonist MPLA induced the strongest local inflammatory reaction with ECM delivery that remained elevated for weeks after injection. Despite MPLA creating an infection-like local environment (**Fig 3**) with SIS ECM, antigen specific cytotoxic function was attenuated compared to the soluble adjuvant alone. This shows that the local scaffold environment can be antagonistic to specific adjuvants and that chronically elevated inflammation alone is not predictive of cytotoxic immunity. Likewise, the cytokine GM-CSF did not convey a substantial benefit to APC recruitment or increase cytotoxic activity with SIS ECM co-delivery despite increasing cellularity. Rather, the most prominent effect of GM-CSF was to increase the acute (1 day) neutrophil response. GM-CSF is often used in combination with other factors to mobilize, attract, and mature APCs at a vaccination site to increase immune recognition in both cell-based and scaffold-assisted vaccine designs alike [16, 17, 34]. ECM scaffolds alone already efficiently attract myeloid cells and promote expression of similar cytokines [10], thus diminishing the role of GM-CSF to potentiate recruitment. This highlights that the same adjuvant may display different activities based on biomaterial type. In contrast to MPLA and GM-CSF, the STING agonist CDA nominally affected the acute inflammatory response and was the most potent cytotoxic inducing adjuvant. CDA was previously shown to be an effective cancer vaccine adjuvant [30, 44, 45], though have yet to achieve that success clinically [46]. CDA is recognized intracellularly by STING expressed by immune and non-immune cells, which then triggers Type I interferon expression to promote T cell survival and expansion [47].

Prolonging vaccine antigen exposure is another mechanism to increase vaccine efficacy in ECM scaffold materials. ECM contains native protein binding motifs that sequester soluble factors (such as cytokines and growth factors) to extend their half-life and enhance activity, and these properties can be preserved in decellularized tissues [48–50]. We found that OVA protein retention was indeed enhanced compared to soluble delivery, possibly via such binding interactions. Conversely, the adjuvants used in the present study have different chemical properties. The cyclic dinucleotide CDA (and analogous cGAMP used for in vivo tracking) is a small molecule that lacks hydrophobic or positively charged domains that may be important to ECM binding and are thus quickly released. The inorganic salt Alum retained both OVA and CDA at the injection site to a greater extent than ECM, though this did not produce a therapeutic benefit suggesting SIS ECM binding affinity was sufficient in the context of vaccination. Further, a substantial proportion of vaccine signal was identified in Alum over 30 days after injection, which is beyond the optimal time scale for T cell priming.

Immune modulation and sustained antigen release may both contribute to the observed synergy between vaccine delivery and the SIS ECM scaffold microenvironment leading to complete regression of established tumors that is not achievable with soluble vaccine components alone. We further established the magnitude of anti-tumor immunity and investigated mediators of the ECM scaffold-assisted vaccine response. Therapeutic cancer vaccination is a more clinically relevant but more challenging model than prophylactic vaccines (delivered before tumor formation) due to the immune suppressive tumor microenvironment that dampens efficacy. Our CTL assay results showed that very low doses of CDA are sufficient to prime OVA antigen specific immunity, yet even a high dose of CDA in a soluble vaccine was ineffective when treating established tumors (**SFig 2B**). Additional cell types or soluble factors may be required to overcome tumor microenvironmental barriers. NK cells are a promising candidate as our cell depletion studies showed that while cytotoxic T cells were strictly required, NK cells may contribute to influence reliability. Ultimately, our SIS ECM scaffold-assisted vaccine formulation exceeded efficacy reported in many prophylactic and therapeutic EG.7-OVA vaccine models [16, 51–53]. Protection from tumor rechallenge several months after vaccination bolsters the importance of adaptive immunity and provides evidence of long-lived memory lymphocyte generation that are positively associated with durable immunotherapy responses in the clinic [54]. SIS ECM scaffold assisted vaccination responsiveness improved with age, which agrees with certain clinical cancer subtypes such as melanoma [38]. In addition to improved response rates in the primary tumor, vaccination also provided partial protection from the parental lymphoma strain EL4 suggesting evidence of epitope-spreading. Additional validation is required but it is plausible that dying EG.7 cells are releasing non-OVA tumor antigens shared with EL4 that are also being presented by APCs to expand the T cell repertoire beyond the vaccine. Epitope spreading has important clinical implications as it overcomes antigen-escape mechanisms of resistance to targeted immunotherapies wherein cancer cell clones downregulating tumor antigen emerge [55].

A key finding of this work is that the pro-healing Type 2-like ECM scaffold immune environment can be used to augment cytotoxic anti-tumor immunity. Traditionally, Type 2 immunity is often considered antagonistic to T cell mediated cancer immunotherapy. For example, Th2 polarized T cells are enriched in breast tumor subsets and correlated with poor outcomes [56]. However, this paradox is tempered by previous studies that demonstrate contextual importance. We previously showed that ECM scaffold immune environments inhibited local melanoma tumor formation *in vivo* via a T cell and macrophage dependent mechanism, providing proof-of-principle that ECM scaffolds can promote anti-tumor immunity [10]. Furthermore, Type 2 cytokines such as IL-4 are pleiotropic and can assist cytolytic immunity in cell-based immunotherapy [57, 58]. Other features of the ECM immune response are more complex such as hybrid M1/M2 macrophage population that is phenotypically distinct from immune suppressive tumor macrophages [10]. Interestingly, the adjuvants we tested did not diminish these Type 2 like signatures, for example *Il4* gene expression in lymph nodes or STAT6 signaling in the scaffold microenvironment, suggesting that these immune states can co-exist. The molecular mechanisms of how biomaterials modulate local immunity *in vivo* is an area of active study, though cellular participants differentiate ECM biomaterials from synthetic polymeric or inorganic materials that drive the chronic foreign body reaction. Persistent neutrophils, Th17 cells, and multinucleate giant cells that accumulate around non-degradable polymers are absent in the ECM response. We showed that both ECM and the inorganic particulate salt Alum enhanced therapeutic tumor regression despite these fundamental differences in immune environment, and additional analyses are necessary to understand whether ECM scaffold specific features are productive or detrimental to immunotherapy.

This study shows that ECM scaffolds prepared from decellularized tissues can enhance cytotoxic T cell priming and improve the efficacy of a therapeutic cancer vaccine when using an appropriate immune adjuvant. Cyclic di-AMP (CDA) induced systemic antigen-specific cytotoxic T cell immunity *in vivo* while not significantly altering immune features that are important to ECM scaffold remodeling. Cytotoxic immunity translated to tumor regression in established tumor microenvironments when used as a therapeutic vaccine. The ECM scaffold immune environment can therefore be synergistic with cancer immunotherapy and is a promising addition to treatment. This expands the toolbox for scaffold-assisted cancer vaccine delivery to include biomaterials that can be applied to promote healing during tissue reconstruction and to merge fields of tissue engineering and cancer immunology.

## Acknowledgements

This research was supported by the Intramural Research Program of the NIH, National Cancer Institute, CCR, Cancer Innovation Laboratory. S.P. and M.T.W. assisted in conceptualization, experimental design, performing experimental procedures, and in writing/editing the manuscript. R.C. assisted with experimental design and performing experimental procedures. I.B. assisted with experimental procedures and data analysis. B.J.H. assisted in experimental design and intellectual feedback. B.N. assisted with animal procedures. The authors thank Dan McVicar, David Wink, Stephen Anderson, Howard Young, Joost Oppenheim, Ji Ming Wang, and Scott Durum of the Cancer Innovation Laboratory for enlightening discussions.

## Methods

### Small intestinal submucosa (SIS) decellularization and cyrogenic milling

Normal porcine small intestine was obtained from Tissue Source LLC (Zionsville, IN) from market weight pigs that were documented as pathogen free (porcine reproductive and respiratory syndrome, porcine epidemic diarrhea virus, porcine delta coronavirus, transmissible gastroenteritis) and complied with ISO 13485. Intestines were flushed of their contents and cut open along its length then mechanically delaminated to remove the muscularis and mucosal layers. The resulting submucosa was cut into 6 inch pieces and decellularized using 4% alcohol/0.1% peracetic acid (v/v, Sigma-Aldrich) and washed thoroughly with 1X PBS and Type 1 water followed by lyophilization and comminuted into an injectable particulate form via cryogenic milling and sieving through 425 µM pore sieve. SIS particles were terminally sterilized via 2×10^6^ rad gamma irradiation on dry ice. Particles were tested as negative for murine viral and bacterial pathogens.

### DNA quantification in decellularized SIS-ECM particles

The DNA was isolated from lyophilized porcine native small intestine (SI) and decellularized SIS ECM particles using DNeasy Blood & Tissue Kit according to manufacturer’s protocol. Briefly, minced lyophilized native SI and SIS ECM particles were digested with proteinase K in ATL buffer at 56°C. Once tissue was completely digested by visual inspection, AL buffer was added and incubated at 56°C for 10 minutes followed by 100% ethanol. The samples were loaded onto DNeasy Mini Spin columns and centrifuged at 8000g for 1 minute. The column was washed with AW1 and AW2 buffer, and DNA was eluted with 200µL of AE buffer with centrifuging at 8000g for 1 minute. Isolated DNA was quantified using Quant-iT PicoGreen dsDNA Assay Kits according to manufacturer’s protocol. Briefly, 100µL of diluted DNA samples were added in triplicate into 96 well plate followed by addition of 100µL of Quant-iT PicoGreen reagent. The plate was incubated for 5 minutes at room temperature in the dark and fluorescence emission at 520 nm quantified after 480 nm excitation using a plate reader (Spectramax i3, Molecular Devices).

### Scanning electron microscopy and SIS particle size quantification

The SIS particles were scattered onto aluminum SEM stubs and sputter coated with either 5 nm thick gold-palladium 30mA for 30second or 4.5 nm thick iridium 30mA for 30 seconds using a K575X sputter coater (EMITECH, Quorum). Images were acquired with Hitachi S-4500 field emission SEM and processed using Quartz PCI (v9) software. Size quantification from 9 separate images with multiple fields of view was manually conducted by blinded observers at the NCI Frederick EM facility (Electron Microscopy Laboratory).

### Mice

Female 7-week old C57Bl/6J mice and SKH-1 hairless mice were obtained from The Jackson Laboratory and housed at the NCI Frederick Laboratory Animal Sciences Program in specific pathogen-free conditions and under 12-hour light/dark cycles. Ethical approval for the animal experiments was provided by the Institutional Animal Care and Use Committee at NCI Frederick (Protocol No. 20-063). Mice acclimated to housing conditions for one week prior to experimental procedures. Mice were euthanized via asphyxiation with carbon dioxide and cervical dislocation.

### Subcutaneous SIS ECM injection and tissue collection

SIS-ECM particles were hydrated with saline to form an injectable suspension. Each SIS ECM dose consisted of 5 mg of SIS particles hydrated with 100 µL of saline for a minimum of 30 minutes and intermittent vortexing. For studies into the effect of immune adjuvant on SIS ECM immune environment, adjuvants were first prepared in saline and then used to hydrate SIS particles as described above for 20 µg CDA, 10 µg MPLA, or 1 µg GM-CSF per SIS ECM dose. For antigen and vaccine studies, 100 µg of OVA protein was included per dose.

SIS ECM particles were subcutaneously injected in the right flank of C57Bl/6 mice for immune response characterization and vaccine studies, or at the tail base for live animal imaging. Briefly, the injection area was shaved and disinfected with alcohol and 100 µL of SIS ECM particle suspension injected (day 0) with or without the described vaccine components. Immune characterization studies were performed 1, 7, or 14 days post implantation. Mice were anesthetized with 2% isoflurane for blood collection via cheek bleeds starting 1 day before SIS ECM implantation (baseline) and then prior to euthanasia at each time point. SIS ECM scaffolds and adjacent subcutaneous tissues were harvested for fixation in 10% neutral buffered formalin for a minimum of 48 hours. Lymph nodes after 7 and 14 days were snap frozen on dry ice for PCR analysis.

### Histologic analysis and multiplex immunofluorescence staining

SIS ECM scaffolds and nearby skin was explanted to ensure that the SIS implant was intact. Explants were carefully trimmed and cut into two halves from the midline over the implant and cut faces were embedded into paraffin for sectioning. Formalin fixed tissues were dehydrated with a graded series of ethanol and xylene for paraffin embedding, sectioned (5 um), and stained for H&E as per standard protocols (Histoserv, Inc.). H&E stained implants were imaged using a Zeiss AxioObserver using a 20X objective (high resolution) or 10X objective tiled images. Immunofluorescence staining and imaging was performed to characterize immune cell infiltrate within SIS ECM scaffolds in combination with vaccine components. For all stains, sections were deparaffinized in xylene followed by rehydration in decreasing concentration of ethanol and then in Type I water. Rehydrated sections were post-fixed with neutral buffered formalin for 15 min followed by PBS wash with Type I water. Antigen retrieval of tissue sections was performed in citrate buffer (pH 6.0) for 20 min at 95-98°C using a steamer and cooled at RT for 20 minutes followed by washing with Type I water. Endogenous peroxidases were quenched by incubating the slides in 3% H2O2 in PBS for 15 minutes. Slides were washed with Type I water and a border was created around the section using PAP pen. Unreacted aldehydes were quenched with 2.24% (0.3M) Glycine (w/v) in TBS-T buffer (Tris Buffered Saline with 0.05% Tween-20) for 5 minutes followed by blocking using 10% BSA in TBS-T for 30 min at room temperature.

Sections were sequentially stained via tyramide signal amplification with full reagent information can be found in Table 1. In brief, each round of staining consisted of incubation with primary the antibody diluted in blocking buffer, 3 washes in TBS-T, incubation with species appropriate HRP-Polymer conjugated anti-IgG secondary for 15 minutes at room temperature (RT), 3 washes in TBS-T, incubation with Opal dye diluted in Opal amplification diluent (Akoya), and 2 water washes. Antibody stripping was performed between rounds using citrate buffer (pH 6.0) for 20 min at 95-98°C. After the final round, slides were counterstained with DAPI (1µg/ml) in PBS for 5 min followed by water washes and cover slipping with fluorescent antifade mounting reagent (DAKO, Agilent).

For myeloid staining, primary antibody labeling conditions and Opal dye pairs were applied in the following order: (1) F4/80 at 1:500 dilution overnight at 4°C and Opal 570 at 1:150 dilution, (2) CD86 at 1:500 dilution for 30 minutes at room temperature and Opal 650 at 1:150 dilution, (3) Ly6G at 1:2000 dilution for 30 minutes at room temperature and Opal 520 at 1:150 dilution.

Minor staining modifications were applied for phospho-STAT staining; antigen retrieval was performed with pH 8.0 Tris-EDTA and PBS buffer steps were replaced with TBS or water. Primary antibody labeling conditions and Opal dye pairs were: (1) pSTAT6 at a 1:200 dilution overnight at 4°C and Opal 570 at a 1:150 dilution, (2) pSTAT1 at a 1:500 dilution overnight at 4°C and Opal 650 at a 1:500 dilution.

### Whole slide imaging and computational pathology

The whole slide fluorescent images were evaluated for the quantification and characterization of immune cell infiltrates immunolabeled for Ly6G, F4/80, and CD86 antibodies. Whole Slide Imaging was performed using an Aperio fluorescent scanner (Leica Biosystems, Wetzlar, Germany) with a 20x objective to detect DAPI, Opal 520, Opal 570, and Opal 650. Image deconvolution, annotations, cell detection, and threshold determination was performed using the opensource QuPath software package (v0.3.3). Annotations were created by a board-certified pathologist to include implant border, interface, and core; the interface was defined as 200 µm concentrically from the border, and the core as greater than 200 µm towards the center. Cell detection was performed using pretrained StarDist convolutional neural networks.

### Quantitative real time PCR of lymph node cytokine expression

The RNA from draining LN (dLN) was isolated using Qiagen RNeasy micro kit according to manufacturer’s protocol. Briefly, the dLN was crushed in liquid nitrogen and 0.5mL Trizol was added to it. 0.1mL of chloroform was added to the sample tube and vortexed vigorously for 15 seconds. The tubes were centrifuged at 12000g for 15 minutes at 4°C and upper aqueous layer was taken out in a 1.5mL tube. 0.25mL isopropyl alcohol was added to the tube and incubated for 10 minutes. The solution was loaded onto RNeasy MinElute spin column and eluted in RNase-free water. The RNA concentration and purity was confirmed using Qubit RNA high sensitivity assay kit and RNA integrity and quality assay Kit. 2.0µg of the isolated RNA was reverse transcribed to cDNA using SuperScript IV VILO master mix (Invitrogen) according to manufacturer’s protocol. The RNA and cDNA were stored at - 80°C till use.

RT PCR was performed in triplicate to quantify the gene expression of Il4 and Ifnγ from cDNA of dLN using TaqMan Gene expression assay. Briefly, the TaqMan gene expression master and TaqMan Assay and cDNA was added in LightCycler 480, 96 well plate. The reaction plate was sealed with adhesive film and centrifuged to collect the contents at the bottom. The sealed plate was run in Roche LightCycler 480 Instrument II and programmed according to the manufacturer’s instruction. The fold change in gene expression was calculated using the 2^-ΔΔCt method.

### Plasma cytokine analysis

The PROCARTAPLEX 21 PLEX kit (Thermo Fisher Scientific, SKU# PPX-21) was used to quantify the various cytokines from mice plasma (Table 2 from SI). The plasma was run by Frederick National Laboratory core facility according to the manufactures protocol on Luminex FLEXMAP 3D instrument and the results were analyzed by Bio-Plex Manager software.

### In vivo cytotoxic T-lymphocyte (CTL) assay

The cytotoxic T-lymphocyte (CTL) assay was performed to assess the functional output of cellular mediated immunity against ovalbumin as a model antigen in vaccinated mice. Briefly, 6-8 weeks old female C57Bl/6 mice were vaccinated with SIS+Adjuvant+ovalbumin (OVA; 100µg) on day 0 followed by a booster on day 7. On day 13, spleens were isolated from naïve C57Bl/6 mice in RPMI media on ice and diced into small pieces and digested using Liberase TL (0.25 mg/ml) and DNAse (0.2 mg/ml) in 5mL RPMI for 15 min at 37°C on shaker. The digestion mixture was grinded through 70µm cell strainer into 50 ml conical using syringe plunger and cold PBS was passed to rinse out cells. Splenocytes were centrifuged at 300g for 5 min at 4°C, and pelleted cells were washed with cold PBS. The splenocytes pellet was resuspended in 5mL 1X RBC lysis buffer for 3 minutes on ice followed by two washes with cold PBS. The splenocytes were divided into two tubes of 10 million cells each. The first tube was labelled with High concentration of Celltracer Violet dye (25µg/mL) and second tube splenocytes were labelled with low concentration of Celltracer Violet dye (2.5µg/mL) by incubating at for 20 mins at 37°C in dark. The unbound dye was quenched by addition of RPMI media with 10% FBS in 1:1 ratio and washed twice with PBS by centrifuging cells at 300g for 5 minutes at 4°C. The splenocytes labelled with high concentration of dye were pulsed with SIINFEKL peptide (2µg/mL) of ovalbumin and low concentration labelled cells were pulsed with scrambled FILKSINE peptide (2µg/mL) of ovalbumin for 30 minutes at 37°C and washed twice with PBS by centrifuging cells at 300g for 5 minutes at 4°C. One million cells were taken from each tube as reference control for analysis by flow cytometry. The cells from two tubes were mixed into 1:1 ratio and intravenously transferred into the previously vaccinated mice (10×10^6^ cells/mice). On day 14, the mice were euthanized and splenocytes were isolated as describe above. The splenocytes were run on the Cytek Aurora Spectral Flow Cytometer followed by analysis on SpectroFlo® software. Specific killing was calculated using the following equation:

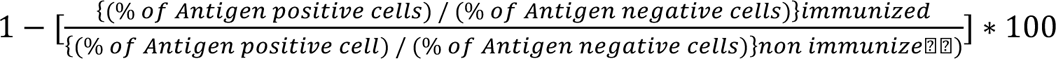

### Ovalbumin fluorescent labeling

Ovalbumin protein was fluorescently labeled for live animal imaging studies using amine reactive IRDye 800-NHS ester (LI-COR). Protein and dye were mixed at a 1:1 equimolar ratio in 2mL 1X PBS (pH 8.4) at RT with continuous mixing for an hour. Labeled protein was purified using Amicon filters (10K MWCO) by centrifugation at 3700g for 10 minutes and washing with 1X PBS. The degree of protein labeling was quantified by absorption at 280nm and 774nm using Nanodrop.

### In-vivo release quantification of OVA and CDA using live animal imaging

We used hairless SKH-1 mice for live animal imaging on IVIS Spectrum In vivo Imaging System (Perkin Elmer). For ovalbumin quantification: The SKH-1 mice were injected at tail base with 100µL of (i) SIS+CDA+OVA, (ii) Saline+OVA-NIR, (iii) Saline+CDA+OVA-NIR, (iv) SIS+OVA-NIR, (v) SIS+CDA+OVA-NIR and (vi) Alum+OVA-NIR and (vii) Alum+CDA+OVA-NIR. The mice were imaged at 5 minutes, 2 hours, 6 hours, daily during 1st week and twice a week for following 3 weeks and once in 4th week. The images were quantified using Living Image software.

For quantifying the CDA analogue cGAMP: The SKH-1 mice were injected at tail base with 100µL of (i) SIS+OVA+CDA, (ii) Saline+OVA+cGAMP-Cy5, (iii) SIS+OVA+cGAMP-Cy5 and (iv) Alum+OVA+cGAMP-Cy5. The mice were imaged at 5 minutes, 2 hours, 6 hours, daily during 1st week and twice a week for following 3 weeks and once in 4th week. The images were quantified using Living Image software.

### Therapeutic vaccination in E.G7-OVA lymphoma tumor bearing C57Bl/6 mice

A therapeutic cancer model was used to evaluate the efficacy of SIS+CDA as cancer vaccine against E.G7-OVA lymphoma tumor model. The E.G7-OVA cell lines are derived from EL4 lymphoma cell line that produces ovalbumin and ovalbumin have been used as model antigen to target cancer cells. Briefly, 6-8 weeks old female C57Bl/6 mice (young mice) and 22-24 weeks old female C57Bl/6 mice (mature mice) were inoculated on the right flank with 0.3 × 10^6^ E.G7-OVA cells and once the tumor volume reached around 75-100 mm^3^ the mice were randomized into different treatment groups followed by a booster dose after 7 days. Group 1 mice were untreated control, Group 2 mice were injected with 100 µL Saline+OVA (100 µg OVA) subcutaneously near the tumor site. Group 3 mice were injected with 100µL of SIS+OVA (5 mg SIS; 100µg OVA) and Group 4 mice received 100µL of Saline+CDA+OVA (20µg CDA; 100µg OVA). Group 5 mice were injected with 100µL of SIS+CDA+OVA (5mg SIS; 20µg CDA; 100µg OVA). We used Alum (Alhydrogel® 2% Aluminium Hydroxide Gel adjuvant) as reference for synthetic material in Group 6 where we injected 100µL of Alum+OVA (100µg OVA) and Group 7, 100µL of Alum+CDA+OVA (20µg CDA; 100µg OVA). The mice tumor volume was measured 3 time a week and monitored for survival till 75 days post primary tumor cell injection. A series of rechallenge experiments were performed on the mice which rejected the tumor and became tumor free. The first rechallenge was given on Day 75, by injecting 0.3X10^6^ E.G7-OVA tumor cells on the contralateral side (left flank) of the primary challenge with an aged matched untreated control mice. The second tumor rechallenge experiment was performed in the tumor free mice 233 days after initial tumor implantation with bilateral injection of 0.3X10^6^ EG.7-OVA tumor cells on one left flank and its parental lymphoma line EL-4 on the right flank. The mice tumor volume was measured 3 time a week and monitored for survival.

### Immune depletion in therapeutically vaccinated with ECM assisted vaccine in E.G7-OVA lymphoma tumor bearing mice

The immune cell depletion was performed to determine the effector immune responsible for anti-tumor immunity. We used the E.G7-OVA tumor model as describe in previous section for this study. We used anti-CD8b antibody to deplete cytotoxic T cells, anti-NK1.1 antibody to deplete cytotoxic NK cells, and IgG antibody was used as isotype negative controls (Table 3 SI).

Briefly, we used 6-8 weeks old female C57Bl/6 mice and inoculated with 0.3X10^6^ E.G7-OVA cells on the right flank and randomized the mice with palpable tumor into different groups and administered two I.V injection of the above mentioned antibodies on day 6 and day 8 post tumor injection and once weekly for the next three weeks to maintain the deletion. Group 1 mice were untreated control, Group 2 mice were injected with 100 µL Anti-IgG antibody (100 µg) intraperitonially (i.p). Group 3 mice were injected with 100µL anti-NK1.1 antibody (100µg) and Group 4 mice received 100µL of anti-CD8b antibody (100µg). Once the tumor volume reached around 75-100 mm^3^ all the mice in from Group 2 to Group 4 were injected with SIS-ECM assisted vaccine, SIS+CDA+OVA (5mg SIS; 20µg CDA; 100µg OVA) subcutaneously near the tumor site on day 9 followed by a booster dose on day 16. The mice tumor volume was measured 3 time a week and monitored for survival.

We also quantified the immune depletion of the three-effector cell types in the blood and spleen after 3 depletions by sacrificing 2-4 mice from each group. We harvested blood and spleen and isolated the splenocytes as described in the CTL assay above. We stained the cells with Zombie NIR to exclude the dead cells, anti-CD45 (BUV395), anti-CD11b (Alexa Fluor 700) to exclude all the myeloid cells, and anti-CD3 (PE) for T cells, anti-CD335 (PE-Cy7) for NK cells, anti-CD4 (Pacific Blue) T cells and anti-CD8a (Alexa Fluor 647) T effector cells (Table 4 SI). Briefly, 1X10^6^ cells from blood and spleen were stained with above mentioned antibody cocktail with addition of Fc block in 100µL FACS buffer (0.5% BSA in 1X PBS). The cells were incubated for 40 minutes on ice and washed with FACS buffer twice by centrifuging at 300g for 5 minutes at 4C. The washed cells were fixed using FIX/Perm buffer (BD Biosciences) for 20 minutes on ice and washed with 1X Perm buffer twice by centrifuging at 350g for 5 minutes at 4C. The cells were finally resuspended in 250µL FACS buffer and acquired on Cytek Aurora Spectral Flow Cytometer followed by analysis on SpectroFlo® software.

