## Supplementary Files for "Extracellular matrix scaffold-assisted tumor vaccines induce tumor regression and long-term immune memory"

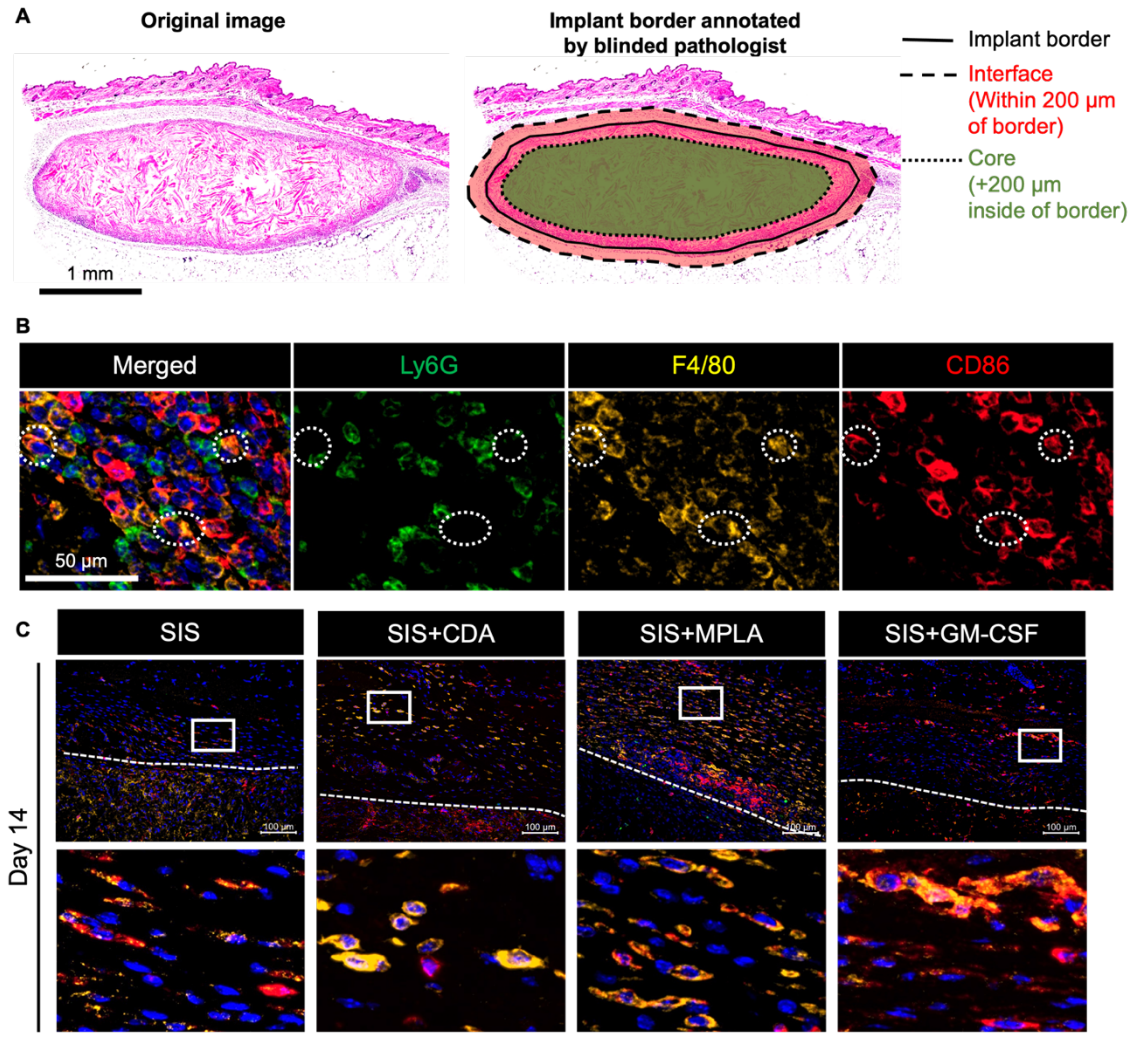


**SFig 1:** (A) H&E images defining the SIS-ECM implant boarder and Interface region and core region, the interface was defined as 200 µm concentrically from the border, and the core as greater than 200 µm towards the center. (B) Multiplex immunofluorescent images depicting the staining of Neutrophils (Ly6G+), macrophages (F4/80+), and antigen presenting cells (APCs, CD86+) on subcutaneously injected SIS ECM particles in C57Bl/6 mice. (C) Multiplex immunofluorescent images of subcutaneously injected SIS ECM particles in C57Bl/6 mice alone or with the immune adjuvants CDA, MPLA or GM-CSF at 14 days post implantation (20X objective). Dashed lines delineate the SIS ECM implant border and boxes detail immune phenotype at this interface.

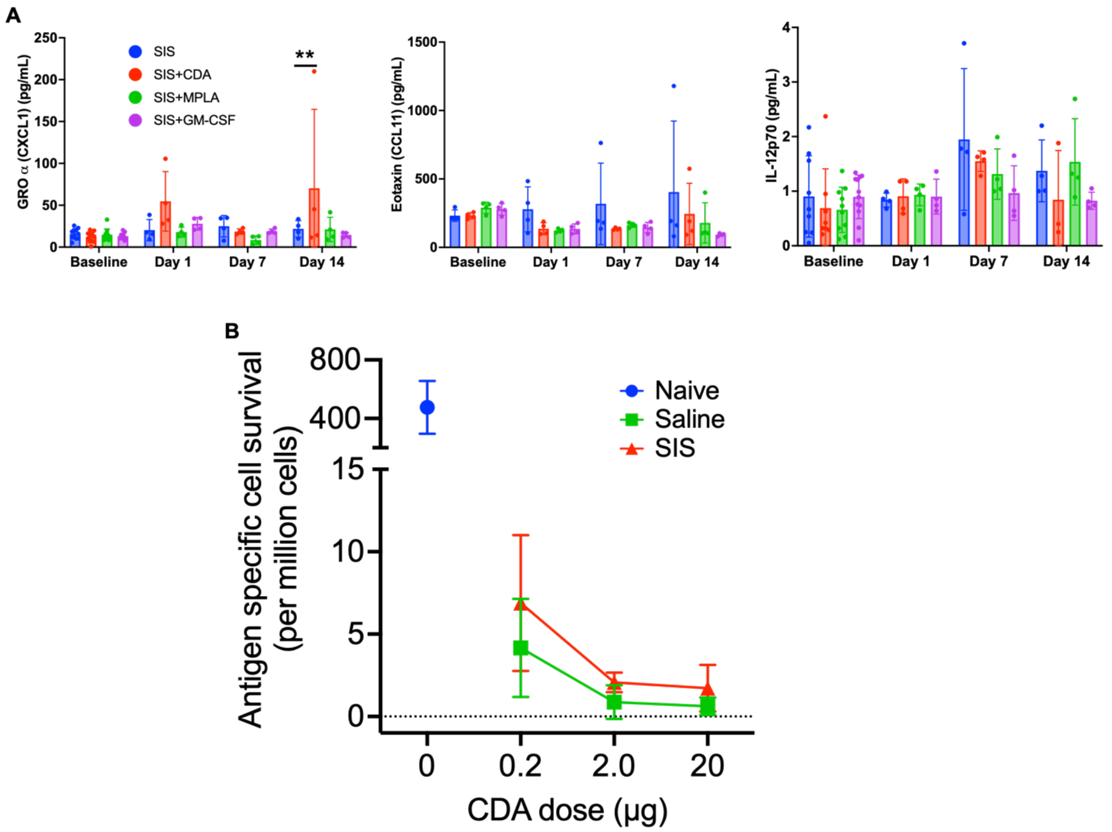


**SFig 2:** Peripheral blood was collected 2 days before SIS ECM implantation (Baseline) then 1, 7, and 14 days post implantation for Luminex analysis. (N=3-4, mean ± SD). **$p < 0.01$, two-way ANOVA with Tukey's multiple comparisons test. (B) Quantification of OVA antigen-specific survival when SIS ECM was co-delivered with different concentrations of CDA. (N=3-4, mean ± SD).

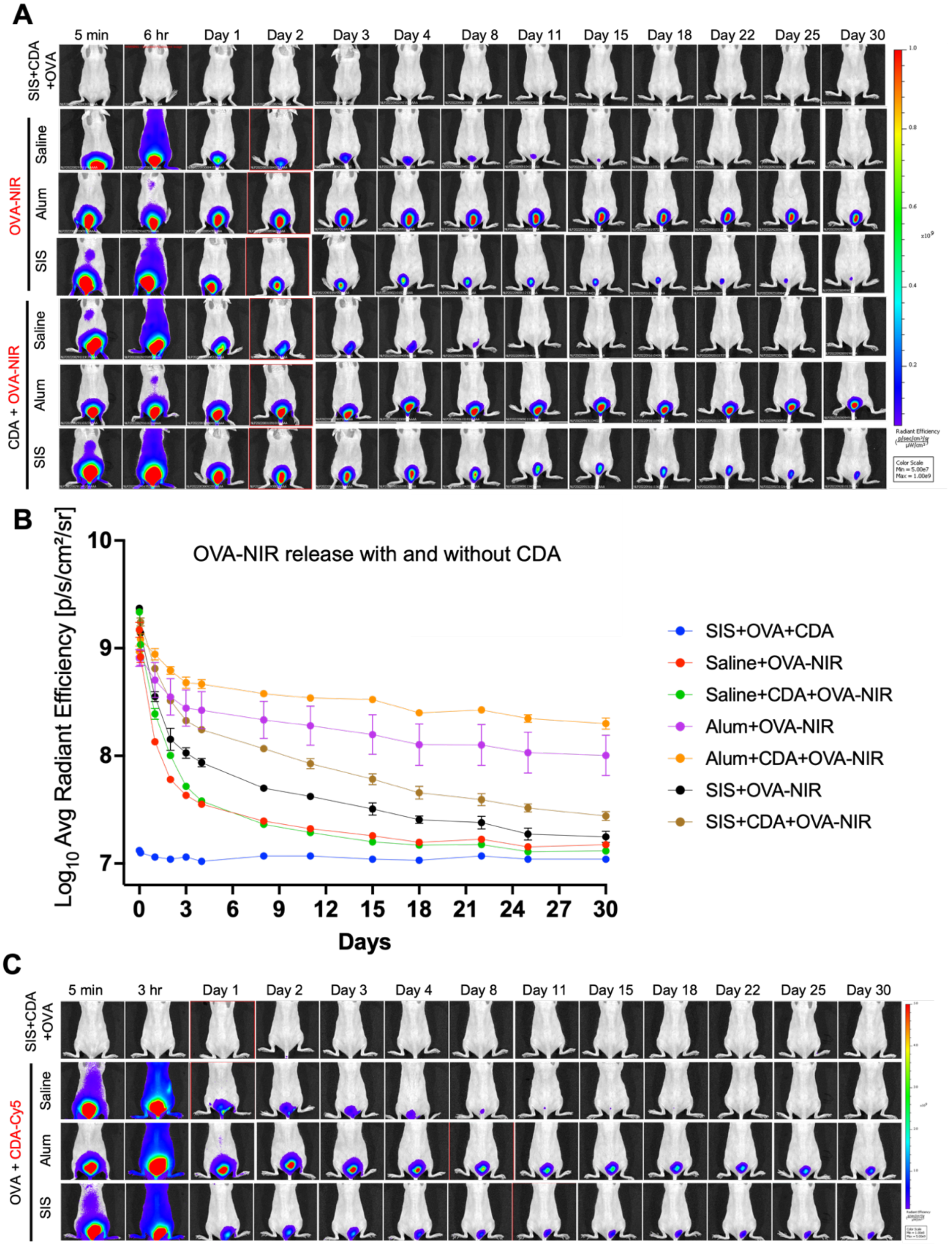


**SFig 3:** (A) Florescent images showing the antigen retention using Licor800 NIR dye conjugated OVA protein when co-delivered subcutaneously with CDA or without CDA and either SIS ECM, Alum, or Saline control at the tail base of hairless immunocompetent SKH1 mice. (B) Fluorescence flux from labeled OVA was quantified at the tail base and normalized to initial signal (5 min post injection) over 30 days. (N=3, mean ± SD) (C) Florescent images showing the cyclic dinucleotide retention using a Cy5 conjugate of the CDA analogue cGAMP co-delivered with unlabeled OVA and either SIS ECM, Alum, or Saline control at the tail base of hairless immunocompetent SKH1 mice.

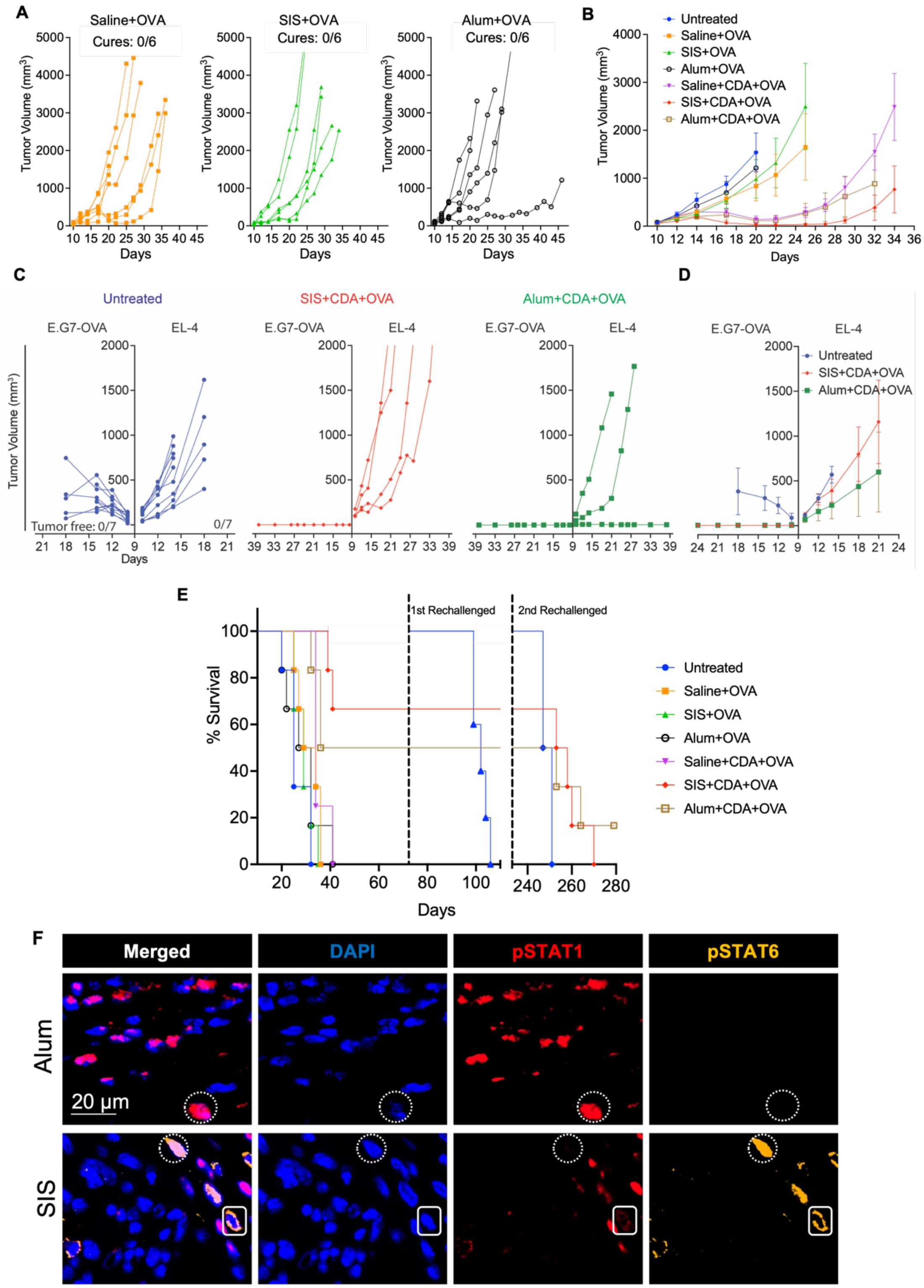


**SFig 4:** (A) Individual tumor growth curves for scaffold assisted therapeutic vaccination, with cures defined as complete and durable tumor regression to a minimum of 75 days. (B) Average tumor growth kinetics (N=6-7, mean ± SEM). (C) Individual tumor growth curves, (D) average tumor growth kinetics, and (E) Overall survival for SIS scaffold assisted therapeutic vaccination upon bilateral rechallenge of E.G-OVA

(Left flank) and EL-4 (Right flank) in rechallenge tumor surviving mice. (F) Multiplex immunofluorescent images showing phospho-STAT1 and phospho-STAT6 staining of subcutaneously injected SIS and Alum in C57Bl/6 mice 7 days post implantation (20X objective).

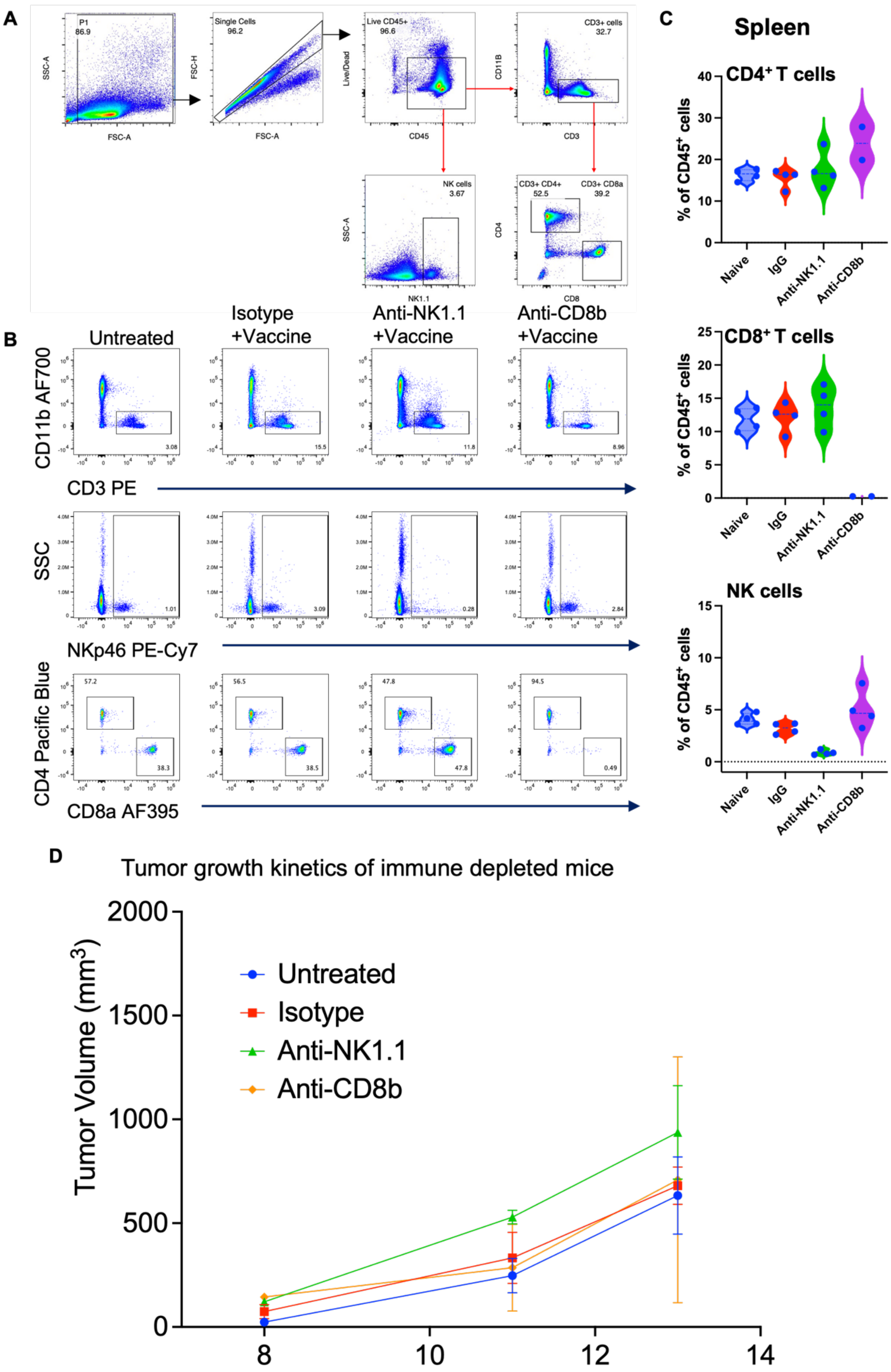

A
P1
86.9
SSC-A
FSC-A
Single Cells
96.2
FSC-H
Live CD45+
96.6
Live/Dead
CD45
CD3+ cells
32.7
CD11B
CD3
NK cells
3.67
SSC-A
NK1.1
CD3+ CD4+
52.5
CD3+ CD8a
39.2
CD4
CD8
B
Untreated
Isotype +Vaccine
Anti-NK1.1 +Vaccine
Anti-CD8b +Vaccine
CD11b AF700
CD3 PE
3.08
15.5
11.8
8.96
SSC
NKp46 PE-Cy7
1.01
3.09
0.28
2.84
CD4 Pacific Blue
CD8a AF395
57.2
38.3
56.5
38.5
47.8
47.8
94.5
0.49
C
Spleen
CD4⁺ T cells
CD8⁺ T cells
NK cells
% of CD45⁺ cells
Naive
IgG
Anti-NK1.1
Anti-CD8b
D
Tumor growth kinetics of immune depleted mice
Tumor Volume (mm³)
Untreated
Isotype
Anti-NK1.1
Anti-CD8b
2000
1500
1000
500
0
8
10
12
14

**SFig 5:** (A) Gating strategy for the quantification of NK cells, CD4 and CD8 T cells. (B) Flow plots showing the gating of CD3, CD4, CD8 T cells and NK cells upon immune cell depletion in different treatment groups. (C) CD8 cytotoxic T cell and NK cell depletion was verified with flow cytometry analysis of peripheral blood. (N=2-4). (D) Average tumor growth kinetics of EG.7-OVA tumor for the immune depleted mice before vaccination with SIS+CDA+OVA.

**Table 1. Histology antibodies**

| Target | Conjugate | Clone | species | Dilution | Company, Cat # |
|---|---|---|---|---|---|
| Ly6G | - | 1A8 | rat | 1:2000 | Biolegend, Cat# 127601 |
| F4/80 | - | BM8 | rat | 1:500 | Biolegend, Cat# 123101 |
| CD86 | - | E5W6H | rabbit | 1:500 | Cell Signaling Technology, Cat# 20018 |
| pSTAT1 | - | 5856 | rabbit | 1:500 | Cell Signaling Technology, Cat# 9167 |
| pSTAT6 | - | D8S97 | rabbit | 1:200 | Cell Signaling Technology, Cat# 56554 |
| Anti-rabbit IgG | HRP polymer | - | - | Neat | Biocare Medical, REF# RMR622H |
| Anti-rat IgG | HRP polymer | - | - | Neat | Biocare Medical, REF# BRR4016H |

**Table 2. List of 21 analytes in Luminex panel (PROCARTAPLEX 21 PLEX (Thermo Fisher Scientific)**

| S.No. | Analyte | S.No. | Analyte | S.No. | Analyte |
|---|---|---|---|---|---|
| 1 | **RANTES (CCL5)** | 8 | IL-1 beta | 15 | IL-10 |
| 2 | **IP-10 (CXCL10)** | 9 | IL-2 | 16 | IFN alpha |
| 3 | **MIP-1 beta** | 10 | IL-22 | 17 | IFN-y |
| 4 | **GRO alpha (CXCL1)** | 11 | IL-3 | 18 | TNF a |
| 5 | **Eotaxin (CCL11)** | 12 | IL-4 | 19 | IL-1 alpha |
| 6 | **IL-12p70** | 13 | IL-6 | 20 | IFN-beta |
| 7 | IL-5 | 14 | IL-17A (CTLA-8) | 21 | GM-CSF |

**Table 3. Immune Cell depletion antibodies**

| Target | Clone | species | Dose (μg/mice) | Company, Cat # |
|---|---|---|---|---|
| NK1.1 | PK136 | mouse | 100 | Bioxcell, Cat# BP0036 |
| CD8β | 53-5.8 | rat | 100 | Bioxcell, Cat# BP0223 |
| Isotype | HRPN | rat | 100 | Bioxcell, Cat# BP0088 |

**Table 4. Flow cytometry reagents**

| S. No. | Target | Conjugate | Clone | species | Dilution | Company, Cat # |
|---|---|---|---|---|---|---|
| 1 | Viability | NIR | - | - | 1:10000 | Biolegend, Cat# 77184 |
| 2 | CD3 | PE | 17A2 | rat | 1:150 | BD, Cat# 100205 |
| 3 | CD45 | BUV395 | 30-F11 | rat | 1:200 | BD, Cat# 565967 |
| 4 | CD4 | Pacific Blue | RM4-4 | rat | 1:200 | BD, Cat# 116007 |
| 5 | CD335 | PE-Cy7 | 29A1.4 | rat | 1:200 | BD, Cat# 137617 |
| 6 | CD8a | Alexa fluor 647 | 53-6.7 | rat | 1:150 | BD, Cat# 100724 |
| 7 | CD11b | Alexa fluor 700 | M1/70 | rat | 1:500 | BD, Cat# 101222 |
